# Spatially expandable fiber-based probes as a multifunctional deep brain interface

**DOI:** 10.1101/2020.10.27.355768

**Authors:** Shan Jiang, Dipan C. Patel, Jongwoon Kim, Shuo Yang, William A. Mills, Yujing Zhang, Kaiwen Wang, Ziang Feng, Sujith Vijayan, Wenjun Cai, Anbo Wang, Yuanyuan Guo, Ian F. Kimbrough, Harald Sontheimer, Xiaoting Jia

**Affiliations:** Bradley Department of Electrical and Computer Engineering, Virginia Tech, Blacksburg, VA; Fralin Biomedical Rsearch Institute, Roanoke, VA; Translational Biology, Medicine, and Health, Virginia Tech, Roanoke, VA; Department of Materials Science and Engineering, Virginia Tech, Blacksburg, VA; Frontier Research Institute of Interdisciplinary Science (FRIS), Tohoku University, Sendai, Japan; School of Neuroscience, Virginia Tech, Blacksburg, VA

**Author notes:** All correspondence and requests for samples should be addressed to X.J.

## Abstract

Understanding the cytoarchitecture and wiring of the brain requires improved methods to record and stimulate large groups of neurons with cellular specificity. This requires miniaturized neural interfaces that integrate into brain tissue without altering its properties. Existing neural interface technologies have been shown to provide high-resolution electrophysiological recording with high signal-to-noise ratio. However, with single implantation, the physical properties of these devices limit their access to one, small brain region. To overcome this limitation, we developed a platform that provides three-dimensional coverage of brain tissue through multisite multifunctional fiber-based neural probes guided in a helical scaffold. Chronic recordings from the spatially expandable fiber probes demonstrate the ability of these fiber probes capturing brain activities with a single-unit resolution for long observation times. Furthermore, using *Thy1-ChR2-YFP* mice we demonstrate the application of our probes in simultaneous recording and optical/chemical modulation of brain activities across distant regions. Similarly, varying electrographic brain activities from different brain regions were detected by our customizable probes in a mouse model of epilepsy, suggesting the potential of using these probes for the investigation of brain disorders such as epilepsy. Ultimately, this technique enables three-dimensional manipulation and mapping of brain activities across distant regions in the deep brain with minimal tissue damage, which can bring new insights for deciphering complex brain functions and dynamics in the near future.

## Introduction

Implantable neural interface devices at a single-unit resolution play an indispensable role in understanding the functional networks in the brain and treating neurological diseases^1–9^. Over the past decades, significant progress has been made in developing highly multiplexed, flexible, and biocompatible neural probes with a single unit resolution for chronic implantation^10–17^. Despite the recent achievement of high resolution and high channel count in these devices^10,18–23^, it remains a major challenge to develop a seamless interface that can map three-dimensional (3D) brain activity across distant regions in the deep brain while minimizing the tissue damage.

At present, indirect methods such as fMRI, MEG, and EEG are utilized to illustrate both the signal pathway across multiple brain regions and the underlying brain circuitry^24,25^. These technologies, however, lack single-unit spatial resolution. Alternatively, optical imaging of genetically encoded calcium and voltage sensors are widely used^26–29^, which are advantageous for mechanistic study in animals but are difficult to translate to humans in the short term. To bridge this technological gap, two-dimensional microECoG and NeuroGrid devices have been developed to enable high-resolution recording across the entire cortical region^18,30,31^. Ultraflexible mesh electrodes have been injected into the deep brain to achieve a long-term stable recording with a minor foreign body response^32–34^, but the recording site is limited to the vicinity of the injection point. Several types of stereo-encephalography electrodes^35^ and multi-shank probes^36–38^ have been developed to probe subcortical tissues, however, the tissue damage arising from multiple implantation sites significantly limits their spatial coverage and their neuroscience applications. A 3D multisite deep brain neural probe with minimal implantation damage would overcome many of these limitations and is described here.

In addition to mapping the electrophysiological signals within the brain, multifunctional neural probes that are capable of modulating the local neural activity provide a powerful technique for studying the brain circuitry^39–43^. Following the development of optogenetics, optical waveguides and micro-LEDs have been integrated into various electrical recording probes to achieve bidirectional neural interfacing^44–52^, which has facilitated the study of brain function relative to behavior. Moreover, localized chemical delivery is another useful method not only for interrupting local brain activities^3^, but also for *in vivo* cell-type identification^53^. To that end, multifunctional fiber-based neural probes have recently been developed using a scalable thermal drawing process, which allows for simultaneous optical stimulation, electrical recording, and drug delivery *in vivo*^54,55^. However, the interfacing sites in these fiber-based neural probes have been restricted to a single location (at the fiber tip) so far, making the broad application of these probes unfeasible. Integrating the optical, chemical, and electrical functionalities in a minimally invasive and 3D probe within the deep brain would provide a versatile and powerful platform for basic neuroscience and clinical applications.

To achieve this goal, we have developed spatially expandable multifunctional fiber-based probes that can interface with neurons using electrical, optical, and chemical modalities simultaneously, while providing a 3D coverage of the deep brain tissue. To create multisite interfaces in each probe (i.e. depth-dependent probes), we expose electrode recording sites, microfluidic channel openings, and waveguide windows at spaced locations along the fiber length using a femtosecond laser micromachining technique. To realize the probe array expansion after implantation, a scaffold with helix hollow channels was used to direct multifunctional fiber probes into brain tissue at specified angles. Chronic recordings validate that these fibers can provide long term neuronal readout with a single-unit resolution at multiple locations. Furthermore, we demonstrate that these 3D multifunctional probes can be used to modulate neural activities in transgenic mice and record distinct brain activities from spaced electrodes. Finally, we are able to detect varying electrographic activities among different brain regions during ictal and interictal periods enabling the detection of seizure foci in a mouse model of chronic epilepsy. Our data suggest that this 3D multiplexed brain interface has the potential to allow for multimodal manipulation and analysis of brain circuitry activity between brain regions under the physiological and pathological state.

## Results

### Depth-dependent multifunctional fibers

In this study, a variety of fiber structures were fabricated using a thermal drawing process (TDP) as previously reported^54–56^. A representative “preform” fabrication process to create a multifunctional fiber-based probe (Fiber S1) is shown in **Fig. 1a**. The waveguide in the center consists of polycarbonate (PC, refractive index n_PC_ = 1.586) and polyvinylidene difluoride (PVDF, refractive index n_PVDF_ = 1.426) as the core and the shell, respectively. Six grooves were machined next to the waveguide, two of which were inserted with BiSn metal alloy as the electrodes while the other four remained open as fluidic channels. To enable a stable drawing process, a sacrificial layer (PC) was added as the outer layer of this preform, which was etched away after fiber drawing. This macroscopic “preform” was then heated and drawn into a ~ 200-meter-long fiber with preserved cross-sectional structure and a 50-300-fold reduction in diameter (**Fig. 1b**). The cross-sectional image of the Fiber S1 is shown in **Supplementary Fig. 1a**. Similarly, Fiber 1 (F1) with one electrode and one microfluidic channel, Fiber 2 (F2) with one electrode, one microfluidic channel, and one waveguide, Fiber 3 (F3) with four electrodes, and Fiber 4 (F4) with four electrodes and one waveguide were fabricated via TDP (**Fig. 1c**), which were used in the following animal experiments. Furthermore, we integrated eight waveguides, four electrodes, and one hollow channel within a single fiber (Fiber S2) using a two-step fiber drawing process, and the cross-sectional image of this Fiber S2 is shown in **Supplementary Fig. 1b**.

**Figure 1:**
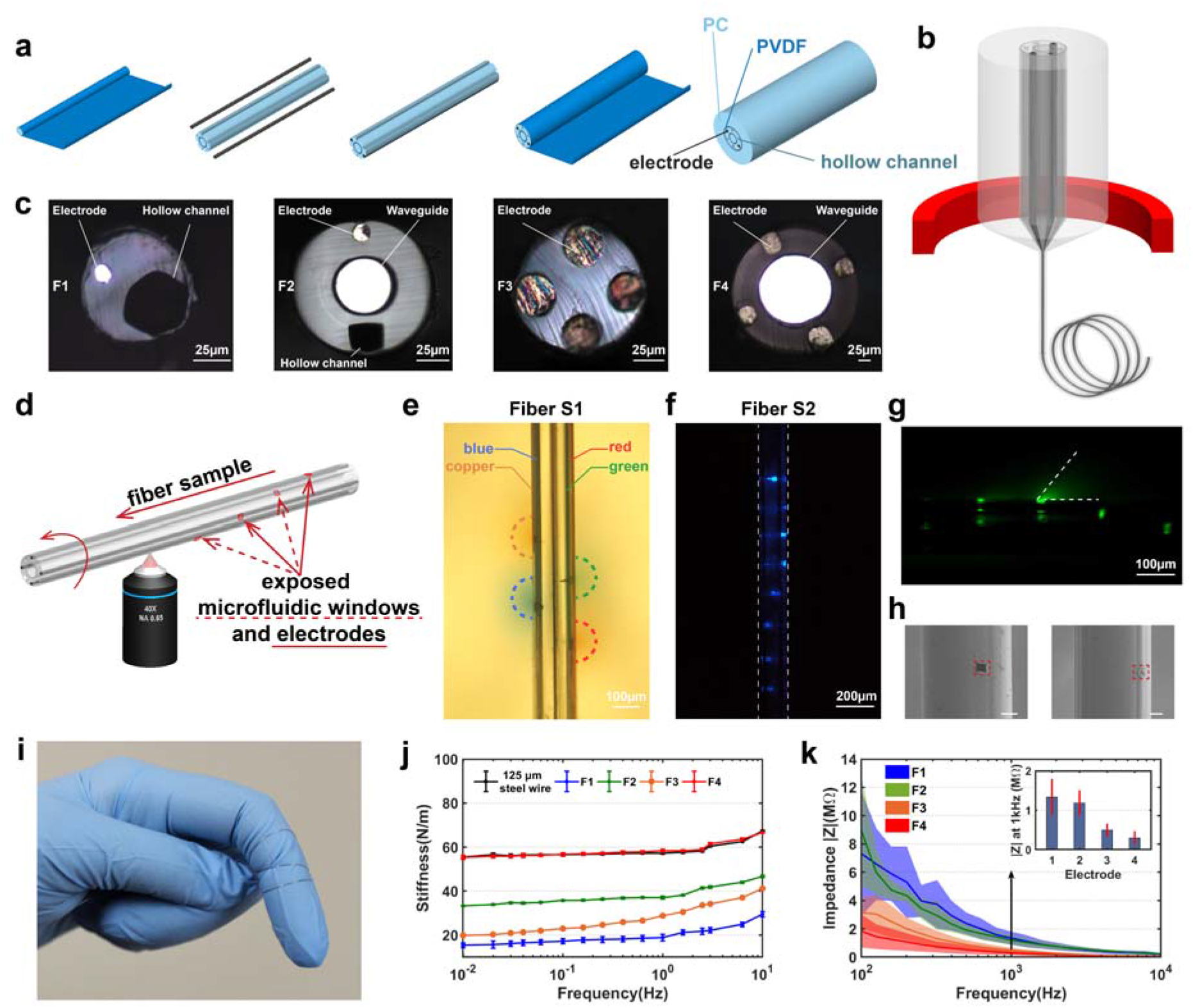
Depth-dependent multifunctional fiber probe. (**a**) A representative multifunctional fiber preform fabrication process (Fiber S1). (**b**) A schematic of the Fiber S1 thermal drawing process. (**c**) Cross-sectional images of fibers used in this study, electrode (BiSn). (**d**) A schematic of femtosecond laser micromachining process on Fiber S1. (**e-h**) Validation of the exposed microfluidic, optical excitation, and electrical windows: e, four microfluidic windows were created on the four hollow channels of Fiber S1 and four different food colors were injected into the four channels respectively while the fiber was embedded in brain phantom; f, an optical image showing the eight optical excitation sites fabricated on the eight waveguides of Fiber S2; g, optical microscope image of the exposed Fiber S2 immersed in a drop of fluorescein excited by a 473 nm laser; h, SEM images of the exposed microfluidic windows and electrodes (scale bar: 50 μm). (**i**) A photograph of functional fibers. (**j**) Bending stiffness measurements of Fiber F1-4. (**k**) Impedance measurements of the BiSn electrodes in Fiber F1-4. All error bars and shaded areas in the figure represent the standard deviation.

To expose multiple optical, electrical, and microfluidic interfacing sites along the fiber length, we used a femtosecond laser micromachining technique. In this technique, a solid-state ultrafast laser based on titanium doped sapphire (Ti: Sapphire) was utilized, which is capable of generating laser pulses of < 10 fs at a wavelength of 800 nm. Our custom-built platform was also capable of rotating the fiber sample during micromachining, enabling 3D patterning along the fiber length (**Fig. 1d**, a more detailed illustration is shown in **Supplementary Fig.1c**). To validate the functionality of the exposed microfluidic and electrical interfacing sites, we created four 20 μm x 20 μm windows on the four microfluidic channels of Fiber S1, respectively, with a 250 μm spacing. This fiber, whose tip was sealed using epoxy to prevent fluidic or electrical leakage at the tip end, was then inserted into a brain phantom (0.6% agarose gel) with four different food colors (1 μl each) injected into these four microfluidic channels, respectively. **Fig. 1e** shows the successful delivery of food colors to four different depths along the fiber.

Beyond exposing the microfluidic channels of Fiber S1, we also created eight 20 μm x 20 μm optical excitation sites with a spacing of 150 μm on the eight waveguides of Fiber S2. We then coupled this micromachined Fiber S2 to a laser source with a wavelength of 473 nm as shown in **Figure 1f.** The light in the optical waveguides is scattered out at the selected locations by ablating partially the cladding and core with the femtosecond micromachining process. Although the light scattering from the eight exposed windows has similar intensity, the directionality in 3D due to their different locations around the fiber circumference causes the windows facing backwards to appear dimmer in the image. In order to have a better understanding of the illumination volumes from these exposed optical excitation sites, we immersed the fiber in a drop of fluorescein solution which emits green light under blue excitation (1% Uranine, Carolina Biological Supply Company). The image shown in **Figure 1g** was taken by an optical microscope and the excitation light was filtered out. Furthermore, we also simulated the light scattering using COMSOL Multiphysics© 5.5 equipped with frequency-domain electromagnetic wave solver to validate the fluorescein image (**Supplementary Figure 2**). The dimension and geometry in the simulation were determined by the actual structure of the fabricated fiber which consists of a waveguide with an 11.5 μm-thick PC core and a 5 μm-thick PMMA cladding. The exposed window is 20 μm x 20 μm in widths and 7.5 μm in depth with ~15° tapering. To mimic the actual surface morphology of a femtosecond laser processed surface, the side and bottom of the exposed window are roughened (R_a_=0.5 μm and 1μm for the side and bottom, respectively). The waveguide core is illuminated evenly with a 473 nm light and the corresponding refractive index of different layers are 1.6023 (PC),1.4976 (PMMA) and 1.3361 (water). From both the fluorescein image and the simulation result, we can observe light scattered from the exposed window with an angle of 50°. There is also some light scattering towards the center of the fiber. However, their intensities are reduced by multiple scatterings and reflections by the electrodes and the hollow channel, and therefore the light intensity emitted from the surface of exposed optical windows is dominant near that area.

Similarly, functional fiber (F3) with four electrodes exposed at a spacing of 50 μm and a size of 20 μm x 20 μm is shown in **Supplementary Fig. 1d**. We further verify the exposed microfluidic window and the exposed electrode by Scanning Electron Microscope (SEM) as shown in **Figure 1h**, respectively. The electrical performance of the exposed electrodes is validated using impedance measurements as shown in **Supplementary Fig. 3**.

We also evaluated the mechanical, electrical, and optical properties of our flexible fiber-based probes. As shown in **Fig. 1i**, the fibers can be easily wrapped around a finger, exhibiting high flexibility. Indeed, most fiber-based probes (F1, F2, and F3) present significantly reduced bending stiffness compared to conventional stainless steel wires (with a diameter of 125 μm) (**Fig. 1j**). Even F4 (at a diameter of 250 μm) shares similar bending stiffness with the steel wire which is much thinner than F4. The impedance of the fiber probes at 1 kHz is found to be 1.3±0.5 MΩ for F1, 1.2±0.3 MΩ for F2, 490±160 KΩ for F3, and 300±170 KΩ for F4, making the fibers good candidates for spike recording (**Fig. 1k**). The optical transmission spectrum was obtained at a wavelength range of 400 – 700 nm, showing broad transmission across the visible range. The transmission loss measured using cut-back method is 0.99±0.03 dB/cm (F2, R^2^=0.99308, n=3) and 1.31±0.03 dB/cm (F4, R^2^=0.99274, n=3) at a wavelength of 473 nm (the excitation peak of channelrhodopsin 2 (ChR2)), indicating that the probes allow sufficient light transmission for optogenetic applications. The bending test at 90° angle and radii of curvature of 1.6-7.9 mm shows no significant bending loss under these deformation conditions (**Supplementary Fig. 4**).

### Spatially expandable multifunctional fiber-based probes

Beyond micromachining to create depth-dependent multifunctional interfaces along a single fiber, helical scaffolding fibers were developed for producing spatially expandable fiber probes. To prepare the preform, four PC tubes (with an inner diameter of 9.53 mm and an outer diameter of 12.7 mm) were bundled together and then wrapped with a thin layer of PC (thickness = 300 μm). After consolidation at 200 °C for ten minutes, four round holes and one square hole were formed in this preform (**Fig. 2a**). Using a customized preform feeding stage that enables simultaneous translational and rotational motion of the preform, we were able to draw helical scaffolding fibers in a controllable manner (**Fig. 2b**). Seven different rotational speeds, 60 r/min, 69 r/min, 93 r/min, 108 r/min, 129 r/min, 147 r/min, and 162 r/min, were employed in this study and the optical images of the side views of the scaffolding fibers drawn at these different rotational speeds are shown in **Fig 2c**. We can observe that as the rotational speed increases, the pitch decreases. **Fig. 2d** demonstrates the relatively linear relationship between the pitch of the scaffolding fiber and the rotational speed, and the average pitch are 23.2 mm, 22.0 mm, 19.3 mm, 15.7 mm, 12.0 mm, 9.0 mm, and 6.7 mm corresponding to each rotational speed, respectively. The cross-sectional geometry of this scaffolding fiber was examined using SEM as shown in **Fig. 2e**. This unique fabrication method also allows us to scale up the numbers of channels in a single scaffolding fiber, such as seven channel-scaffolding fiber as shown in **Fig. 2f**.

**Figure 2:**
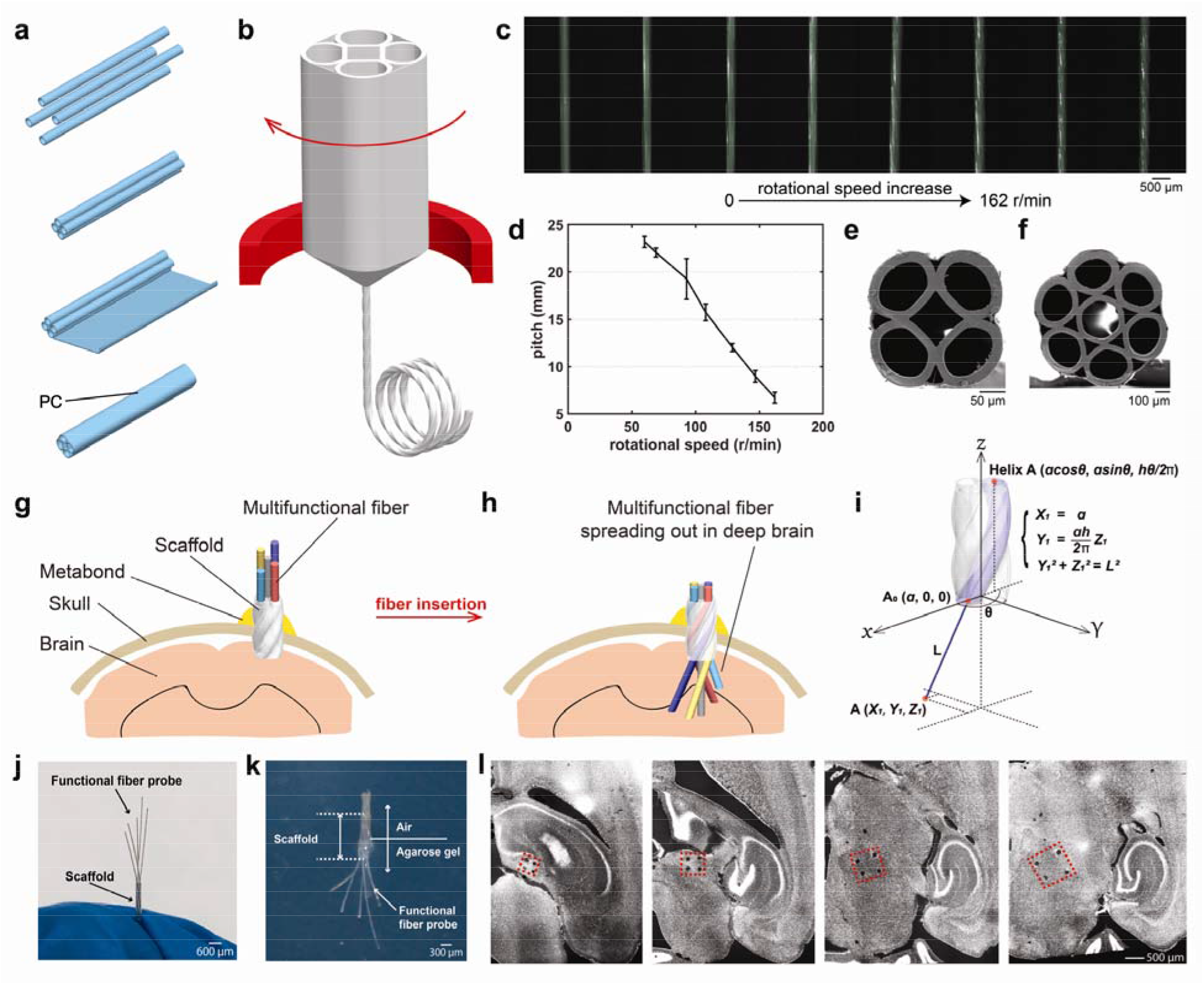
Spatially expandable multifunctional fiber-based probes. (**a**) Scaffolding fiber preform fabrication process. (**b**) Fiber drawing process with customized rotational feeding stage. (**c**) Side-view optical images of the scaffolding fibers drawn at different rotational speeds. (**d**) Plot showing the relatively linear relationship between the pitch and the rotational speed utilized in this study. (**e, f**) SEM images of the scaffolding fibers with 5 and 7 hollow channels. (**g-i**) Schematics of the employment of the spatially expandable functional fiber probes: g, scaffolding fiber is inserted into the brain and affixed by Metabond^®^; h, functional fiber probes are further inserted into the brain tissue through the scaffolding fiber; i, mathematical model of the locations of the inserted fiber probes. (**j**) Pre-validation of the expansion of the inserted functional fiber probes before implant surgeries. (**k**) Validation of the expandable fiber probes in the brain phantom. (**l**), Transverse DAPI-stained brain slices from the mouse with the spatially expandable fiber probes implanted for one week (n=10). Representative images are shown taken from 10 animals. All error bars in the figure represent the standard deviation.

After the scaffolding fiber is fabricated, an array of multifunctional fibers was inserted into the hollow channels of the scaffold. The scaffolding fiber is used to guide individual functional fibers as well as to alter the direction of the inserted fibers when exiting the scaffold. As illustrated in **Fig. 2(g-h)**, the employment of the spatially expandable fiber probes involves two steps during the implant surgery. First, we lower the scaffold to a predetermined depth of the brain and apply Metabond^®^ to fix the scaffolding fiber on the skull (**Fig. 2g**). Second, we further extrude the multifunctional fiber arrays through the scaffold by a calculated length until they reach their targeted locations (**Fig. 2h**) and fix the whole device onto the skull using a dental cement. The penetration depth of the fiber is controlled by the depth of the scaffolding fiber, the extrusion length of the inserted fiber, as well as the extrusion angle of the scaffold. The location of the inserted fiber can be further confirmed by the equation shown in **Fig. 2i**. For the fiber inserted into the center hole, it will stay straight when exiting the scaffold. The center point of the helix bottom is set to be the origin of the coordinates. The scaffolding fiber (**Fig. 2e**) comprises four helixes that intersect with the XY plane at (*a*, 0, 0), (0, *a*, 0), (−*a*, 0, 0), and (0, - *a*, 0), respectively. For the Helix A (*acosθ, asinθ, hθ/2π*), *a* is the radius, θ is the twisting angle of the scaffolding fiber, and *h* is the pitch of this circular helix. *L* is the length of the inserted fiber that is in direct contact with brain tissue. The point of the inserted fiber that intersects with the XY plane is defined as A_0_ (*a*, 0, 0) and the endpoint of the inserted fiber is defined as A (*X_1_, Y_1_, Z_1_*). As the inserted functional fiber that exits the scaffold will naturally form a tangent line to the Helix A, *X_1_* equals *a*. Based on a given value of *a, L*, θ, and *h*, the coordinates of the endpoint of the inserted fiber A (*X_1_, Y_1_, Z_1_*) can be calculated by two θ-independent equations: 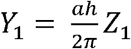 and *Y*_1_^2^ + *Z*_1_^2^ = *L*^2^. Similarly, the locations of the inserted fiber that comes from Helix B (−*asinθ, acosθ, hθ/2π*), Helix C (−*acosθ, −asinθ, hθ/2π*), and Helix D (*asinθ, −acosθ, hθ/2π*) that intersects with XY plane at B (0, *a*, 0), C (−*a*, 0, 0), and D (0, −*a*, 0) are 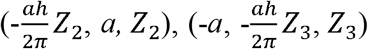, and 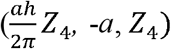, respectively. Further calculation is needed for transferring the coordinates to the commonly used ones relative to bregma depending on how the scaffolding fiber is placed. As these coordinates suggest, we can observe a pattern that the diagonal lines of these four points are perpendicular to each other in the XY plane and the intersect point of these two diagonal lines would determine where the scaffolding fiber should be placed in the XY plane.

Before implanting the probes into animals, we will first insert all the functional fiber probes into the channels of the scaffolding fiber individually and move the scaffolding fiber up and down to confirm the expansion of the functional fibers as shown in **Fig. 2j**. To visualize the spatial expansion of fiber probes through the scaffolding fiber, we inserted the scaffolding fiber to a brain phantom (0.6% agarose gel), followed by the further insertion of the functional fiber probes as shown in **Fig. 2k**. To further confirm the location of the expanded fiber probes, we implanted the scaffolding fiber and functional fiber probes into the thalamus region of the mouse brain (n = 10) for one week and stained the transverse brain slices (50 μm thickness) for cell nuclei using DAPI. We can clearly observe the damage caused by the scaffolding fiber as well as the functional fiber probes from the implants as shown in **Figure 2l**. We can also track the trace of the expanded fiber probes and observe that the distance between the inserted functional fiber probes increases from slice to slice. The square shape formed by the four damage points rotated evenly from brain slice to slice which fits the expectation of linear insertion of the functional fiber probes when exiting the scaffolding fiber. Based on the damage points from the stained brain slices, we found that the expanded fiber probes can reach the targeted region with an average success rate of 81.3% per site (number of successful sites/number of total sites) and 60% per animal (number of successfully targeted animals/number of animals).

As the previous results showed, the angle of the inserted functional fiber can be further tuned by adjusting the rotational speed during the thermal drawing of the scaffolding fiber, which indicates that the scaffolding fiber can guide the functional fiber into arbitrary locations in the deep brain. Furthermore, utilizing scaffolding fiber with more hollow channels and combining it with depth-dependent probes would allow us to sample dense brain regions in three dimensions.

### Spike recording and burst-suppression recording using spatially expandable fiber probes

To evaluate the functionality of the spatially expandable fiber probes in recording localized brain activities with certain distances, we implanted fiber probes with 5 individual branches (Fiber F1) into the different parts of the hippocampus of wild type mice (n=5) as schematized in **Fig. 3a**. **Fig. 3b** shows a mouse with these fiber probes implanted for 5 weeks. The recording was conducted in these mice under continuous anesthesia induced and maintained by isoflurane. Two out of the five implanted fiber probes detected spike activities 5 weeks post-implantation (**Fig. 3c-h**). The electrophysiological signals recorded 5 weeks after implantation were bandpass-filtered to examine the local field potential (LFP, 0.3-300 Hz) and spiking activity (0.3-5 kHz). On one electrode, we found one distinct cluster of spikes *via* principal component analysis (PCA). This spike activity is most likely axonal, as indicated by the sharp voltage drop and short duration waveforms (**Fig. 3c**)^57^ and its peak and repolarization voltages are shown in **Fig. 3d**. The other electrodes captured spiking activity from multiple neurons, from which two spike clusters can be well isolated *via* PCA (**Fig. 3e-h**), where the quality of the isolation was assessed by L-ratios and isolation distance (0.0084 and 147.5, respectively). Thus, based on the recording obtained from these two spaced electrodes, we could detect very different activity patterns with our spatially expandable fiber probes providing useful insights for future investigation of brain functions.

**Figure 3:**
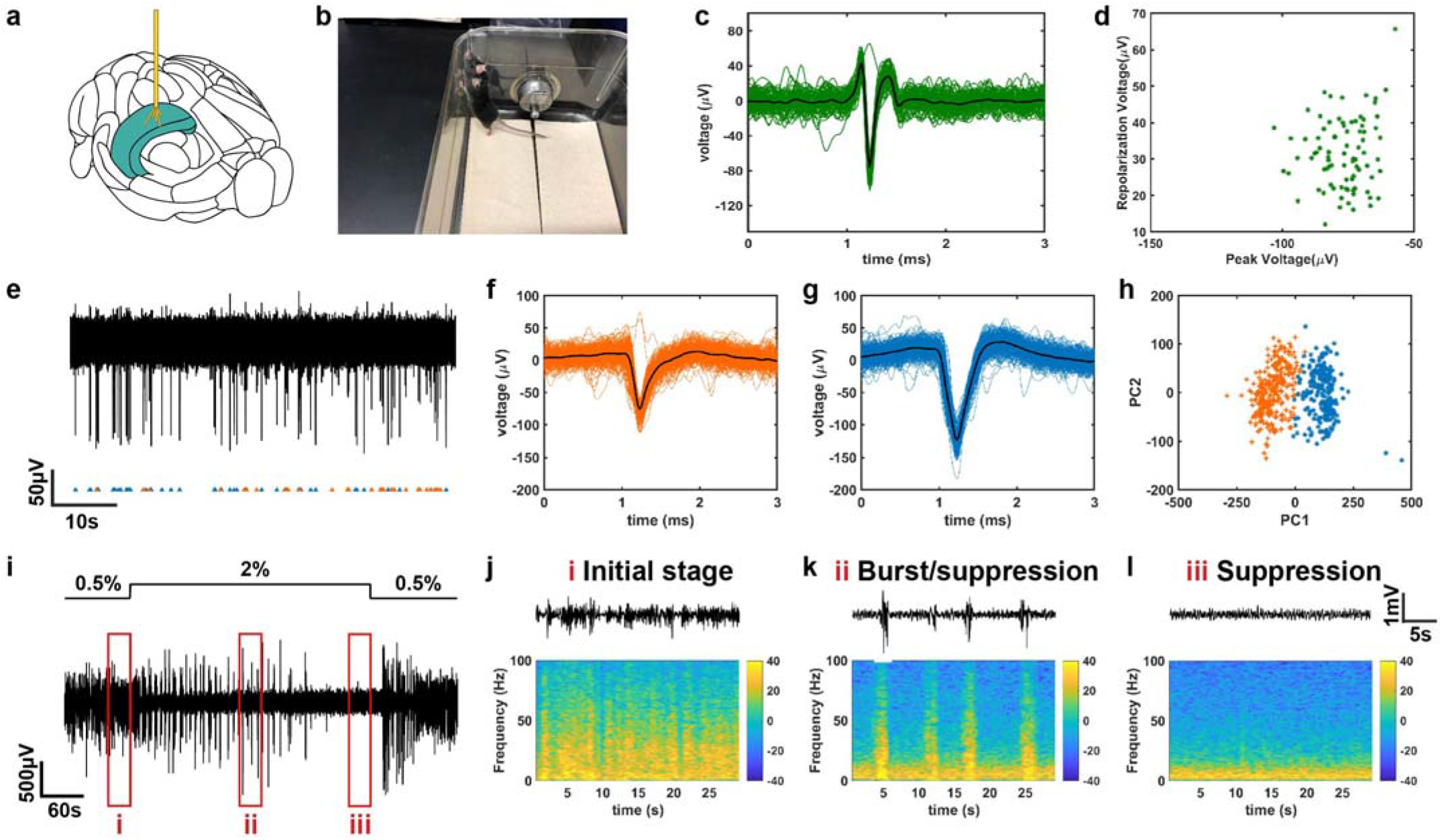
Spontaneous activity investigation by spatially-expanded functional fiber probes (F1, n=5). (**a**) An illustration of the spatially-expanded fiber probe targeting the hippocampus region in a mouse brain. (**b**) A photograph of a wild type mouse with small electrodes implanted for 5 weeks. (**c**) A single cluster segregated 5 weeks after implantation from electrode one and (**d**) it’s peak voltage and repolarization voltage. (**e-h**) Filtered endogenous activity 5 weeks after implantation from electrode two containing two separable units. (f-g) Action-potential shapes of the two units. (h) Principal-component analysis (PCA) of the two units. (i) Local filed potentials recordings 8 weeks after implantation from electrode one with the alternating anesthesia level (0.5-2% v/v isoflurane). (**j-l**) Different levels of LFP activity recorded under the varying concentration of isoflurane, where (j) shows the initial stage (before the increase of isoflurane level), (k) shows burst/suppression stage (transition period) and (l) shows suppression stage (deep anesthesia period). Representative images are shown taken from 5 animals.

We also altered the isoflurane levels during recording (0.5-2% v/v) and observed characteristic time and dosage-dependent changes in the LFP (**Fig. 3i-l**) 8 weeks post-implantation. **Fig. 3j** depicts the power spectrum of LFP obtained under a lower level of anesthesia (0.5% v/v isoflurane). After the concentration of isoflurane was adjusted to 2% v/v, we observed burst suppression (**Fig. 3k**), a hyperexcitable brain state induced by gas anesthetics where alternating high voltage activities (burst) and flatline (suppression) periods appear quasiperiodically^58,59^ (**Fig. 3j**). When the animal was in deep anesthesia, a general suppression of the LFP occurred (**Fig. 3l).** Both respiration rate and responsiveness to toe pinch decreased as the anesthetic level was increased. During deep anesthesia (i.e., suppression), there was a loss of toe pinch withdrawal. The LFP activity returned to the level as measured during the initial stage of recording once the concentration of isoflurane was decreased to 0.5% v/v. Four electrodes out of the five implanted fiber probes presented similar results during the administration of isoflurane (**Supplementary Fig. 5**)

### Optogenetic control and electrophysiological readout with implanted fibers

In this study, we first exposed Fiber F4’s four electrodes via femtosecond micromachining with a spacing (distance between neighboring electrodes) of 0.5 mm, 1.1 mm, and 0.9 mm from top to bottom. The waveguide in the fiber probe was coupled to a silica optical fiber using a direct ferrule-to-ferrule coupling, which was connected to a laser source with a wavelength of 473 nm. The electrodes were connected to the headstage of the recording system via pin-connectors for electrical readout. These depth-dependent fiber probes were implanted into *Thy1-ChR2-YFP* mice, with four electrodes designed to target the cortex, the hippocampus, and two places in the thalamus, and the optical stimulation was delivered at the bottom of this fiber probe (n=5, **Fig. 4a**). Laser pulses at frequencies of 10 Hz, 20 Hz, and 100 Hz with a pulse width of 5 ms and a power density of 7.3 mW/mm^2^ were applied and all the four electrodes detected optically evoked neural activities corresponding to 10 Hz laser pulses 5 months after implantation (**Fig. 4b**). After normalization of the peak-to-peak value of the optically excited signals, we observed that electrode four which was the closest to the stimulation site detected optically excited signals with the highest amplitude systematically (**Supplementary Fig. 6a**). We also conducted the same stimulation and recording experiments in a perfused brain (**Supplementary Fig. 7**) confirming the signals we detected were not artifacts.

**Figure 4:**
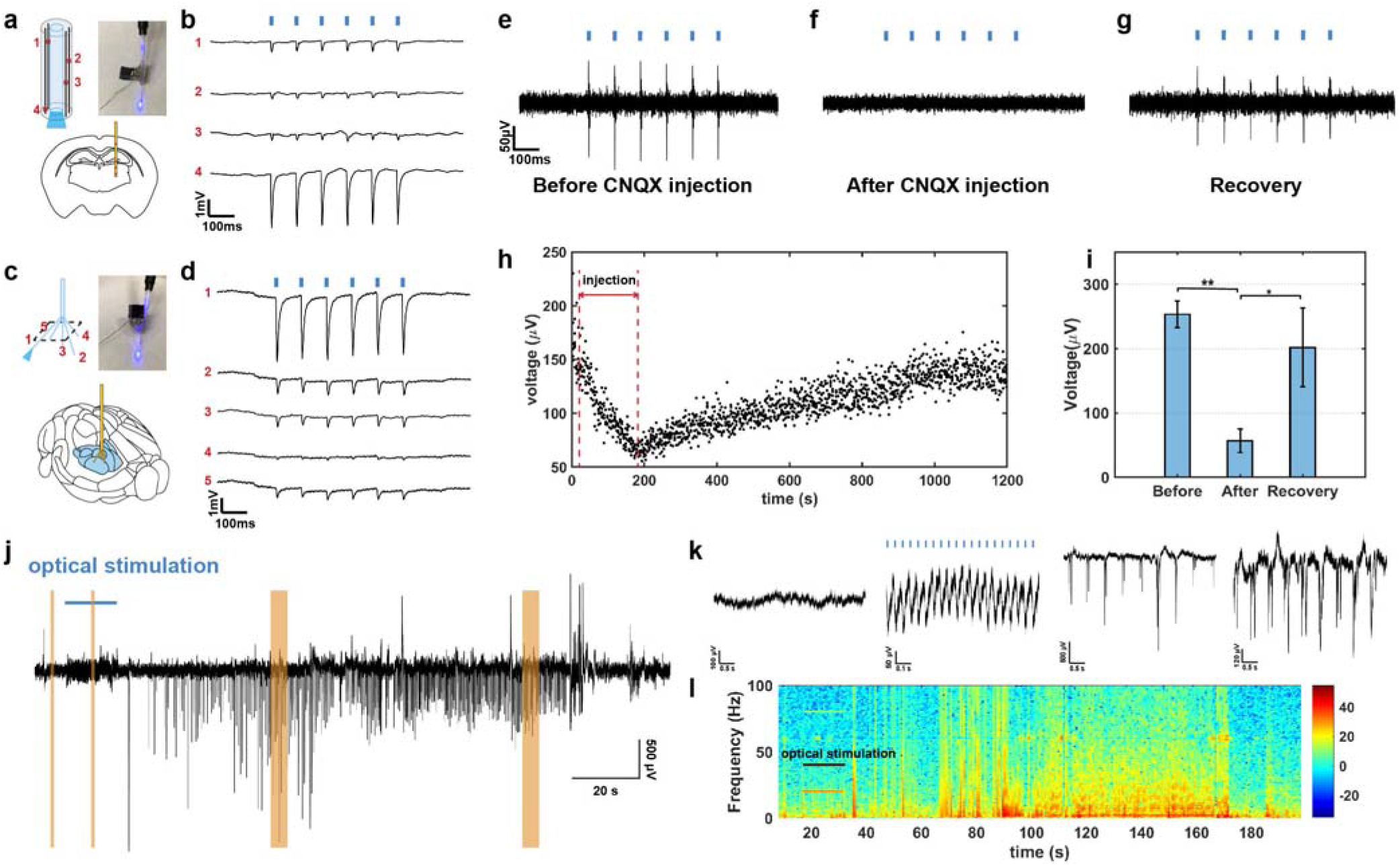
Simultaneous multisite optogenetic stimulation, electrical recording, and drug delivery. (**a**) An illustration of the depth-dependent multimodal fiber probe targeting cortex, hippocampus and thalamus regions in a transgenic Thy1-ChR2-YFP mouse brain and a photograph of the assembled device coupled to the 473 nm laser. (**b**) Unfiltered electrophysiological recording from depth-electrodes 1-4 during optogenetic stimulation (10 Hz, 5ms pulse width, 7.3 mW/mm^2^) 5 months after implantation (n=5). (**c**) An illustration of the spatially-expanded multimodal fiber probe targeting the thalamus region in a transgenic Thy1-ChR2-YFP mouse brain and a photograph of the assembled device coupled to the 473 nm laser. (**d**) Unfiltered electrophysiological recording from electrodes 1-5 during optogenetic stimulation (10 Hz, 5ms pulse width, 7.3 mW/mm^2^) 3 months after implantation (n=5). (**e-g**) Filtered (0.3 - 5 kHz) electrophysiological recording from a F2 multifunctional probe (e) before, (f) 30 s after and (g) 1180 s after CNQX administration through the multifunctional fiber during optogenetic stimulation (10 Hz, 5ms pulse width, 7.3 mW/mm^2^). (**h**) Peak-to-peak amplitude values of each detected optically evoked spiking activity show a decrease in amplitude induced by CNQX administration and a gradual recovery phase after the termination of CNQX delivery. (**i**) Comparison of the mean peak-to-peak amplitude values measured before, during and, after CNQX injection (Repeated measure one-way ANOVA Tukey’s multiple comparisons test, **p<0.01 (Before vs. After, p=0.0044), *p<0.05 (After vs. Recovery, p=0.0134), n=3). (**j**) Seizure-like afterdischarges were induced by repetitive optical stimulation (20 Hz, 5ms pulse width, 7.3 mW/mm^2^) using spatially expandable fiber probes (n=10). (**k**) Zoomed in details of the recording results highlighted in Fig. 4j. (**l**) Power spectrum density plot of the representative trace shown in Fig.4j. Representative images are shown taken from ≥ 3 animals analyzed per each experimental group. All error bars in the figure represent the standard deviation.

We also performed optical stimulation at different brain regions along the depth-dependent fibers and recorded distinct brain activities from spaced electrodes. To our knowledge, both the size of the optical window and the light intensity will have effects on the optically evoked signal. Thus, we have exposed the optical windows with the same sizes on the side of the fiber probes at different depth to ensure the same excitation area and similar light intensity. In this specific experiment, we exposed the four electrodes of the Fiber F4 with a spacing of 0.5 mm, 0.8 mm, 1.2 mm from top to bottom to target the brain regions of cortex, CA1, CA3, and thalamus. In the meantime, the optical window is exposed on the side of the fiber right next to the top (cortex, n=6), middle (CA1, n=5) or bottom (thalamus, n=6) electrode to elucidate the neural activity from the same brain region with different optical excitation sites. The measured light efficiency from the laser source to the exposed window is 0.67% (**Supplementary Figure 8**), and the power density in our experiments is 9.1 mW/mm^2^. To block the light leakage from the bottom tip of the fiber, we applied sufficient black epoxy to cover the tip of the fiber probes. We found that the electrode closest to the optical excitation site detected the optically evoked signal with the highest amplitude (**Supplementary Fig. 9**), consistent with our other optical stimulation and recording results (**Fig. 4b**). Furthermore, we employed spatially expandable fiber probes (five F2 probes with a scaffolding fiber) to the brain region of the thalamus and only fiber one was connected to the laser source while the others were simply connected for electrical readout (n=5, **Fig. 4c**). Similarly, laser pulses with a pulse width of 5 ms, a power density of 7.3 mW/mm^2^, and frequencies of 10 Hz, 20 Hz, and 100 Hz were delivered to the thalamus region right next to electrode one. All five fiber probes detected optically excited neural activities with 10 Hz laser pulses 3 months post implantation (**Fig. 4d**) and electrode one observed the highest peak-to-peak amplitude with normalization (**Supplementary Fig. 6b**).

To assess the chemical delivery function in conjunction with the electrical recording and optical stimulation capabilities, the F2 probe’s hollow channel was connected to a miniaturized tubing and implanted to the thalamus region of *Thy1-ChR2-YFP* mice. 2.5 μl of the synaptic blocker 6-cyano-7-nitroquinoxaline-2,3-dione (CNQX, 0.1 mM, 80nl/s) was injected intracranially to the thalamus region through a microfluidic channel of the F2 probe (n=3). We bandpass-filtered the signals from 300 to 5000 Hz to study the activities of optically stimulated neurons. Before CNQX injection, optically evoked neural activities (10 Hz, 5-ms pulse width, 7.3 mW/mm^2^) were observed consistently (**Fig. 4e**). After CNQX injection, we observed no responsive neural activities corresponding to laser pulses (**Fig. 4f**) and neurons started to respond to laser pulses after several minutes of recovery (**Fig. 4g**). We calculated the peak-to-peak amplitude of each optically evoked neural activity during the entire experiment and we can observe the gradual decrease and increase of the amplitude which represents the drug intervention and recovery processes, respectively (**Fig. 4h**). We also observed a reduced peak-to-peak amplitude of the recovered signals compared to those before drug injection (**Fig. 4i**: repeated measure one-way ANOVA Tukey’s multiple comparisons test, **p<0.01 (Before vs. After, p=0.0044), *p<0.05 (After vs. Recovery, p=0.0134, n=3). The data shown in **Fig. 4e-i** were bandpass-filtered in the frequency range of 0.3-5 kHz while the unfiltered raw data still presents optically stimulated signal before and after CNQX injection (**Supplementary Fig. 10**), which indicates that CNQX only affected neurons close to the microfluidic channel while the neighboring neurons that were not in contact with CNQX still responded to laser pulses.

To further demonstrate the multifunctional capability of the fiber probes, we have also employed the spatially expandable fiber probes (fiber F2) for seizure induction via repetitive optical stimulation and simultaneous electrical recording. It has been previously reported that repetitive optical stimulation can perturb the normal hippocampal activity to induce seizure-like afterdischarges^60^. Here, we implanted the spatially expandable fiber probes (Fiber F2) to target the brain regions of cortex and hippocampus (rostral CA1, caudal CA1, and CA3) and performed repetitive optical stimulation to one location (rostral CA1) to induce seizure-like afterdischarges (n=10). During the stimulation, both 10 Hz and 20 Hz light pulses were delivered for 10-30 seconds at a pulse width of 5-15 ms and multiple optical stimulation trials were performed. We can observe that large spikes appeared following optical stimulation and gradually became synchronous in all these four areas (**Fig. 4j-l** and **Supplementary Figure 11**). These seizure-like afterdischarges continued for a duration of 7.5-131.5 s without any optical stimulation. These results show that our multifunctional spatially expandable fiber probes allow for the manipulation and recording of neural activities across multiple brain regions, which can be useful for both basic neuroscience and disease applications.

### Multisite in vivo recordings by fiber probes in a mouse model of infection-induced epilepsy

We also investigated the utility of depth-dependent fiber probes and spatially expandable fiber probes for studying dynamic spatiotemporal structure of seizure using a mouse model of viral infection-induced temporal lobe epilepsy. In this model, wild type mice infected intracortically with Theiler’s murine encephalomyelitis virus (TMEV) show acute behavioral seizures during the first week of infection, exhibit clinically relevant pathological and physiological changes in the hippocampus, survived the infection and later develop chronic epilepsy after approximately two months of infection^61,62^. Detection of seizures in this model has been reported previously by video encephalography (vEEG) studies that used traditional stainless steel electrodes implanted either epidurally^63^ or in the dentate gyrus^61^. These studies were limited by their use of single-channel recording, and thus, could not provide information on the brain regions implicated in seizure generation and propagation. Our multisite fiber-based probes could be very useful to detect network activity from various regions while minimizing damage to the brain.

We implanted depth-dependent fiber probes in mice either vertically (F3) to the skull (straight implant) for targeting cortex (CTX), CA1 and CA3 areas of hippocampus, and hypothalamus (HY) (straight implant, n=3), or at an angle (F3) to the skull (angular implant, n=4) for targeting CTX, CA1, CA3, and amygdala (AMG) (**Fig. 5b and 5f**). The spatially expandable fiber probes were implanted to target CA1, CA3, and thalamus **(Fig. 5j)**. The CA1 region of the hippocampus is highly susceptible to TMEV-induced cell death with a partial or complete loss of pyramidal cell layer, whereas the pyramidal neurons in the area CA3 remain largely intact during acute seizures^64^. The neuronal circuitry in the CA3 region becomes hyperexcitable during epileptogenesis following TMEV infection^65^. Therefore, two electrodes of the fiber probe were inserted into the CA1 and CA3 regions to detect any difference in the field potential in these two regions. The remaining electrodes were targeted to cortex, amygdala, and hypothalamus to detect generalized electrographic activity during the propagation of seizures. **Fig. 5a** shows an example of EEG recording obtained by a depth-dependent fiber probe implanted in the amygdala of TMEV-infected mouse. Various electrographic features associated with seizures such as preictal spiking, the gradual evolution of amplitude and frequency of spikes during the ictal phase, post-ictal spiking and post-ictal suppression were detected by the fiber probe. Representative EEG traces from each brain region during a convulsive seizure (refer to Supplementary Movie 1 for seizure video and EEG recording) and their corresponding frequency distributions are shown in **Supplementary Fig. 12** (**straight implant**) and in **Fig. 5(g-h)** (**angled implant**). Similarly, **Fig. 5(c-d)** shows EEG traces and their corresponding frequency distributions for a nonconvulsive seizure (**straight implant**). The power spectrum of electrographic discharges during the ictal period was computed to compare seizure severity between the brain regions tested (**Fig. 5e and 5i, Supplementary Fig. 12)**. For a straight implant, mean powers of signals recorded from CTX, CA3, and HY were almost identical for convulsive seizures, whereas cortex showed significantly higher activity during nonconvulsive seizures compared to other brain regions (**Fig. 5e**: CTX vs. CA1 – *p<0.05, **p<0.01, ***p<0.001, ****p<0.0001; CTX vs. CA3 – ^‡^p<0.05, ^‡‡^p<0.01; CTX vs. HY – #p<0.05, ##p<0.01; n=14 seizures). For angular implant, mean powers of signals detected by electrodes implanted in CA3 and amygdala were significantly higher than signal intensity in cortex and CA1 (**Fig. 5i**: CA1 vs. CA3 and CA1 vs. AMG – ^††††^p<0.0001; CTX vs. AMG – #p<0.05, ##p<0.01; n=20 seizures). The mean power of signal recorded from CA1 was significantly lower than that from other brain regions for convulsive seizures detected by both straight and angled implants, which reflects TMEV-induced neuronal damage predominantly in the CA1 region (**Fig. 5l and 5m**). Similarly, the spatially expandable fiber probes also detected robust electrographic activity in CA3 compared to CA1 (**Fig. 5k**, n=12 seizures, n=3 animals).

**Figure 5:**
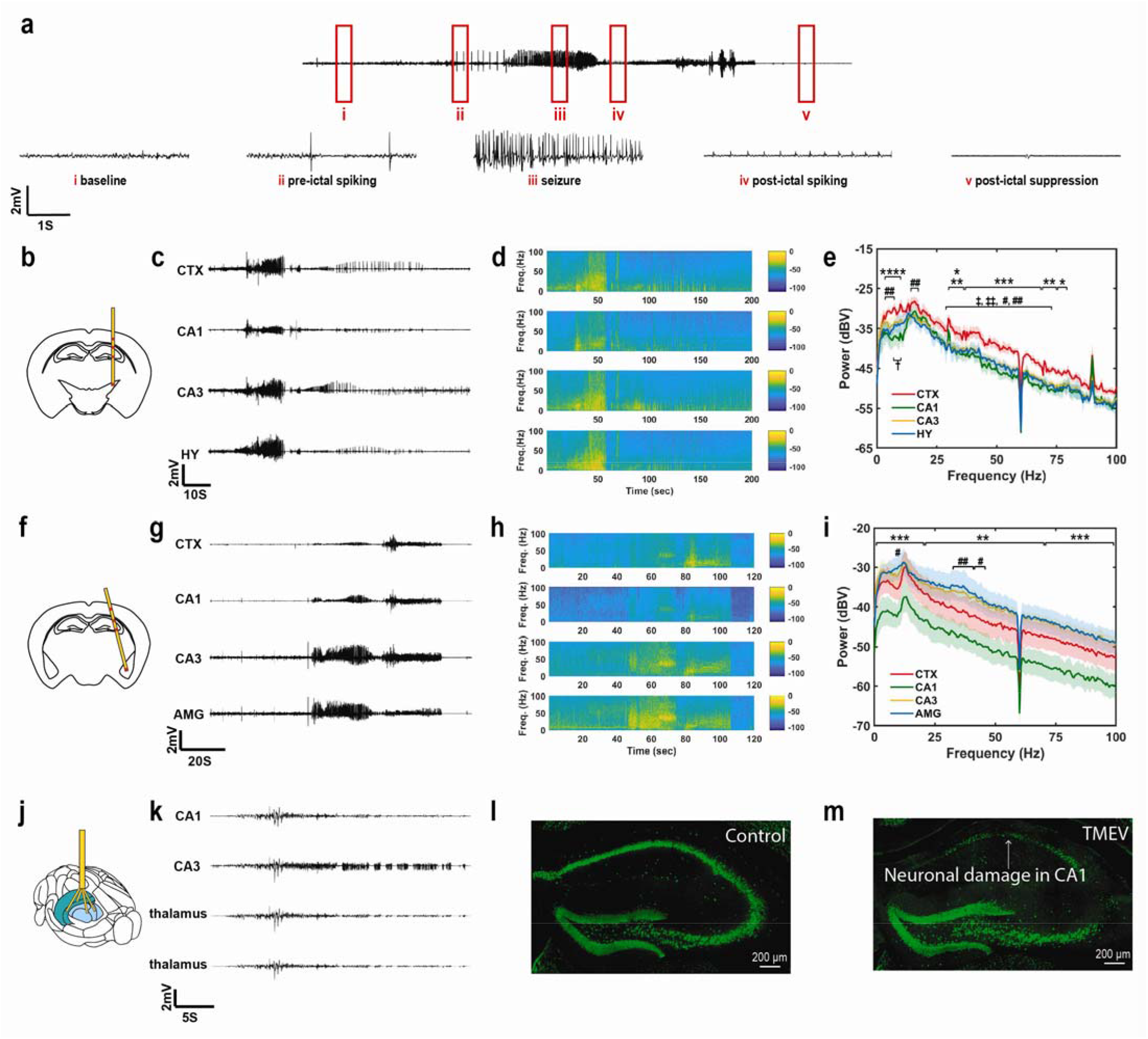
Multisite recording of seizure activity in various brain regions in a mouse model of infection-induced epilepsy. (**a**) An example of electrographic recording by a fiber probe implanted in the amygdala of a TMEV-infected mouse is shown. Various stages of mouse behavior before, during and after convulsive seizure are identified in red rectangles and the corresponding magnified traces are depicted below: baseline (non-ambulatory physiological stage), pre-ictal spiking (periodic electrographic discharges before or during seizure initiation), seizure (high frequency spikings during fully generalized tonic-clonic seizure), post-ictal spiking (periodic electrographic discharges immediately after seizure), and post-ictal suppression (behavioral arrest following convulsive seizure). (**b**) Diagram of coronal section of a mouse brain shows the location of depth-dependent fiber probes in cortex (CTX), CA1 and CA3 regions of the hippocampus, and hypothalamus (HY) as indicated by red marks along with the probe assembly (straight implant, n=3). (**c-d**) Representative traces of electrographic recordings from CTX, CA1, CA3, and HY during a nonconvulsive seizure and their corresponding frequency distribution (straight implant). (**e**) Power spectrum density plot for non-convulsive seizures show that mean power of signal recorded from CTX was significantly higher compared to mean power from other brain regions for various frequency bands between 1-100 Hz (Two-way ANOVA, Tukey’s multiple comparisons test; CTX vs. CA1 – *p<0.05, **p<0.01, ***p<0.001, ****p<0.0001; CTX vs. CA3 – ‡p<0.05, ‡‡p<0.01; CTX vs. HY – #p<0.05, ##p<0.01; n=14 seizures). (**f**) Diagram of the coronal section of a mouse brain shows the location of depth-dependent fiber probes in the cortex (CTX), CA1, CA3, and amygdala (AMG) as indicated by red marks along with the probe assembly (angular implant, n=4). (**g-h**) Representative traces of electrographic recordings from CTX, CA1, CA3, and AMG during a convulsive seizure and their corresponding frequency distribution (angular implant). (**i**) Power spectrum density plot for convulsive seizures show significantly higher mean power of signal recorded from CA3, AMG and CTX compared to that in CA1 between 1-100 Hz (Two-way ANOVA, Tukey’s multiple comparisons test; CA1 vs. CA3 and CA1 vs. AMG – ††††p<0.0001; CA1 vs. CTX – **p<0.01, ***p<0.001; n=20 seizures). (**J**) Diagram of a mouse brain showing the location of two spatially expandable fiber probes implanted in CA1 and CA3 regions of the hippocampus and the other two in the thalamus (n=3). (**K**) Similar to depth-dependent fiber probes, representative traces of electrographic recordings from the hippocampus and thalamus obtained by spatially expandable fiber probes indicate higher network activity in the CA3 region. (**l-m**) Immunohistochemical staining of nuclear neuronal protein (NeuN, in green) in brain slices shows the neuronal loss in the CA1 region of the TMEV-infected mouse with seizures (m) compared to the sham-injected control mice (l). The loss of neurons in the CA1 region may explain the lower power of electrographic activity detected by fiber probes in CA1. Representative images are shown taken from ≥ 3 animals analyzed for each experimental group and were chosen to reflect the mean value of quantitative data. All shaded areas in the figure represent the standard deviation.

Both clinical and animal studies have identified a crucial role of the amygdala, a part of the temporal lobe, in epileptogenesis^66^. A large number of patients with temporal lobe epilepsy (TLE) shows extensive neuropathology in the amygdala, in addition to hippocampal damage. Animal studies suggest that the amygdala is even more susceptible to the generation of seizures than the hippocampus, which may explain faster development of electrical kindling-induced seizures in the amygdala than in the hippocampus. In addition, anxiety and depression are common comorbidities associated with TLE, and the neuronal circuitry in the amygdala and hippocampus is strongly associated with the pathophysiology of anxiety and depression^67^. Indeed, TMEV-infected epileptic mice also show anxiety behavior. Therefore, severe electrographic activity detected in the amygdala and the CA3 region of the hippocampus during TMEV-induced convulsive seizures provides a basis for further investigation of changes in network connectivity between the amygdala and hippocampus in this model.

### Evaluation of fiber probe and stainless steel microwire biocompatibility using immunohistochemistry

Reactive tissue response to our multifunctional fiber-based probe was assessed using immunohistochemical analysis of surrounding brain tissue from mice implanted for four-weeks with either a multifunctional probe Fiber F1 or a conventional stainless steel probe (n=3). The presence of glial fibrillary acidic protein (GFAP) was used to assess astrocyte reactivity to the probe, ionized calcium-binding adaptor molecule 1 (Iba1) was used as a marker of microglial response, and lastly, the neuron-specific protein NeuN was used to analyze neuronal density. Representative images from the multifunctional probe and stainless steel probe are presented in **Figure 6**. Neuronal density, microglia density, and astrocyte reactivity were compared between all groups, and no significant difference was observed, as shown in **Figure 6e-g**. We also validated the biocompatibility of the Fiber F2 by implanting the Fiber F2 probe or stainless steel wire into the hippocampus region of wild type mice for four weeks (n=3) as shown in **Supplementary Fig. 13**.

**Figure 6:**
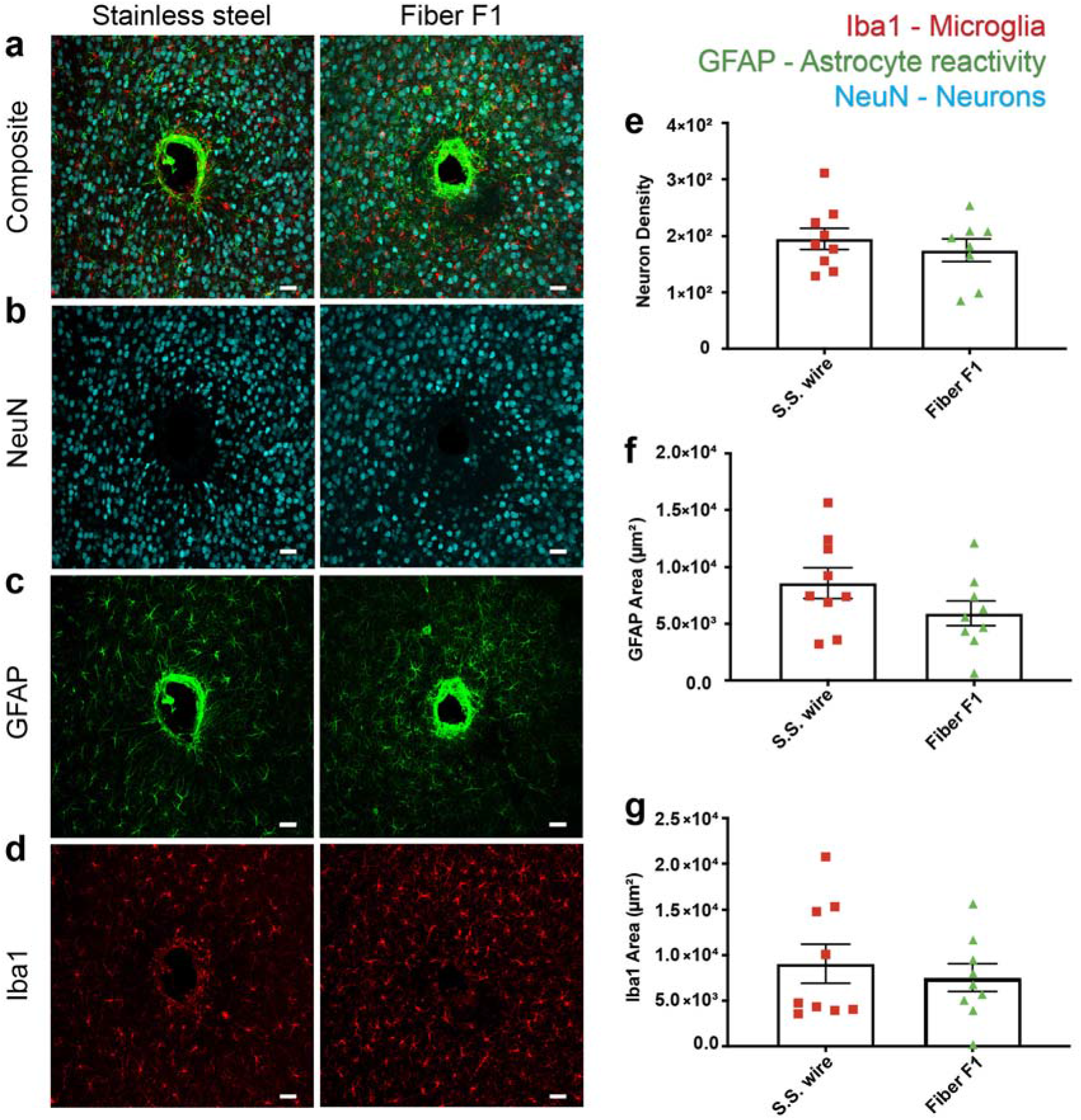
Biocompatibility study of the functional fiber probes (Fiber F1) and stainless steel wire. (**a-d**) Immunohistochemical comparison of tissue reaction to chronically implanted Fiber F1 (n=3) and conventional stainless steel wire (n=3) after four-week implantation. Neurons were labeled with NeuN (cyan), astrocytes with GFAP (green), and microglia with Iba1 (red). (Maximum intensity projections of 50 optical sections covering entire slice thickness taken at the implantation site). (**b, e**) Neuron density, calculated by counting NeuN labeled neurons, was not significantly different between the groups. (**c, f**) Astrocyte and microglia (**d, g**) reactivity, measured as the area of GFAP- or Iba1-positive cells respectively, were not significantly different between groups. Significance was determined by student’s t-test. Error bars on bar graphs reflect the standard deviation. Two-to-three brain slices from each mouse (n=3) per experimental group were used for immunohistochemical analysis. Representative images were chosen to reflect the mean value of quantitative data. (Scale bar: 40 μm)

## Discussion

Here we presented the development and application of spatially expandable multifunctional fiberbased probes for mapping and modulating brain activities across distant regions in the deep brain. The multisite multifunctional interfaces along a single fiber are created using a femtosecond laser micromachining technique. Furthermore, the integration of a helix scaffolding fiber with multifunctional fiber arrays fully extends the capabilities of fiber-based neural probes from a 1D single-site interface to 3D multi-site brain interfaces. These probes provide a powerful platform for recording localized spike activities and field potentials and performing optogenetic and chemical modulation of the activities in spaced deep brain regions. Our customizable fiber probe is also readily compatible with commercial EEG setup allowing us to detect multiple foci and different brain activities among brain regions of interest in the TMEV-infected seizure model. Recordings in epileptic mice illustrate the power of the spatially expandable fiber probes to simultaneously record from distant brain regions allowing circuit analysis of the epileptic brain. In the future, a higher density of electrodes and optical or chemical interfacing sites in these spatially expandable fiberbased probes can be explored to enable a larger recording or stimulation sample. Closed-loop systems that combine the detection of seizure foci with localized intervention can be developed which can significantly improve the effective treatment of seizure before clinical onset. Finally, these probes can enable a leap forward towards the understanding the brain circuitry in the hard-to-access deep subcortical regions, and facilitate the development of new therapeutic methods for treating various brain diseases, including epilepsy, Parkinson’s disease, addiction, depression, autism, etc.

## AUTHOR CONTRIBUTIONS

S.J., D.P., X.J., H.S., I.K., S.V., and Y.G. designed the study. S.J. fabricated all the fibers included in the study. S.J. obtained the impedance spectra. Y.Z., S.J, and Z.F. evaluated optical properties. K.W. and W.C. performed bending stiffness measurements. S.Y., A.W., Y.G., and S.J. conducted femtosecond laser micromachining on functional fiber probes. S.J. connected multifunctional fiber probes. S.J. performed *in vivo* implantation surgeries and collected electrophysiological data. D.P., H.S., and S.J. conducted TMEV injections and collected seizure related data. J.K. wrote the MATLAB script and analyzed recording data. I.K., W.M., S.J., and H.S. performed immunohistochemistry. All the authors contributed to the writing of the manuscript.

## ACKNOWLEDGEMENTS

X.J. gratefully acknowledges funding support from National Science Foundation (ECCS-1847436).

H.S. gratefully acknowledges funding support from US National Institutes of Health (R01-NS036692, R01-NS082851 and R01-NS052634).

## Methods

### Multifunction fiber fabrication

Many fiber structures have been prepared in this study. They all start from the preform fabrication which we described in detail below. After the preform fabrication, the macroscopic preform was put into a customized furnace. With controlled temperature and stress, the preform was drawn into thin fibers. The temperature we used during fiber drawing are 140 °C (top), 285 °C (middle), and 110 °C (bottom).

For fibers used as spatially expandable probes to detect seizure activity (F1), two rectangular grooves were machined on the PC rod, one of them was filled by a BiSn electrode while the other one remained hollow as a fluidic channel. Then the pre-structured rod was wrapped with a few layers of PVDF and an additional PC layer which subsequently formed a preform after the consolidation process.

For spatially expandable fiber probes with optical stimulation capability (F2), a thin layer of PVDF and a relatively thicker layer of PC were added to a PC rod. Two grooves were then machined on this thicker PC layer while one of them was filled by the BiSn electrode. Two layers of PVDF and PC were subsequently wrapped around the rod and put in the oven for the consolidation process.

For fibers with four electrodes to probe for the seizure activities in wild-type mice (F3), four rectangular grooves were machined on a PC rod and all of them were filled with BiSn electrodes. Then a few layers of PVDF and PC were wrapped around the structure and then consolidated.

For fibers used in the *Thy1-ChR2-YFP* mice (F4), we first rolled five layers of PVDF film (Mcmaster) and polycarbonate films (laminated plastics) onto a PC rod (Mcmaster), which subsequently formed a solid rod after the consolidation process in vacuum at 200 °C. Next, we machined four rectangular grooves (2mm x 2mm) on the solid PC layer which were filled by the BiSn (Indium Corporate) electrodes. Then, a few layers of PVDF were rolled onto the electrode-filled rod to form the insulating layer, which was then followed by additional layers of PC for the stable thermal drawing process.

For fibers used in the demonstration of a multifunctional fiber with multisite drug delivery capability (Fiber S1), a thin layer of PVDF followed by layers of PC films were first rolled onto a PC rod. After the consolidation of the films, six grooves were machined on the outer PC layer and two of them were filled with the BiSn electrodes. Then additional PVDF and PC layers were added to the rod to form the complete preform structure.

For fibers with eight waveguides, four electrodes, and one hollow channel (Fiber S2) a two-step drawing process was used. We first drew a rectangular waveguide fiber consisting of a PC core and a PMMA cladding. Then twelve grooves were machined on a PC tube (fabricated by wrapping PC films onto a Teflon rod and consolidating then), and four of them were inserted with CPE (conductive polyethylene) electrodes while the other eight were filled with the waveguide fibers. After that, additional PVDF and PC layers were wrapped onto the PC rod and consolidated.

For spatially expandable fiber probes fabrication, obtaining proper scaffolding fiber is crucial. This was achieved by putting four or seven PC tubes together with a thin layer of PC films as protection. Taking the advantages of the rotating motor on our customized fiber drawing tower, the fiber formed a helix structure due to the rotational degree of the rotating motor. The rotating motor was controlled by a motor and the rotational speed varies corresponding to the voltage applied to the motor. The voltage used in this study include 0, 3.3 V, 4.1 V, 5.0 V, 5.7 V, 6.5 V, 7.2 V, and 8.0 V, resulting in rotational speed of 0, 60 r/min, 69 r/min, 93 r/min, 108 r/min, 129 r/min, 147 r/min, and 162 r/min, respectively.

### Electrochemical Impedance Spectrum

First, fiber probes of two centimeters were prepared and the inner electrodes were electrically connected to the copper wire (connecting details in Neuro Probes Assembly). The impedance Spectrum results were acquired via a potentiostat (Interface 1010E, Gamry Instruments). During the measurements, two-electrode experiments were performed with fiber probes as a working electrode, Pt wire (Basi) as counter electrode and reference electrode and 1x phosphate-buffered saline (PBS, Thermo Fisher) as electrolyte by an AC voltage of 10mV (10 Hz – 100 kHz).

### Stiffness measurements

The stiffness test was carried out using a dynamic mechanical analyzer (DMA Q800, TA Instruments) installed with the single cantilever module. The tested fiber length is 10 mm, which equals to the distance between the fixed and movable clamps. The vibrating magnitude is 20 μm. We tested the stiffness of the fibers together with steel wire as the comparison group under different frequencies ranging from 0.01Hz to 10 Hz. Three repeated measurements of each sample sets were taken at a temperature of 37°C.

### Optical properties characterization

To obtain the optical transmission spectra, we connected a halogen light source (DH1000, Ocean Optics) to the optical waveguide using the ferrule-to-ferrule coupling method. The transmitted light was collected at the other side of the fiber with a fiber-coupled spectrometer (Flame, 200-1,025nm, Ocean Optics). The transmission spectra are normalized by a light source spectrum. To quantify the transmission loss, the fiber probe ferrule end was coupled to a silica fiber patchcord by a mating sleeve (Thorlabs) which was air-coupled to a DPSS laser (Laserglow Technologies, 100 mW maximum power, wavelength λ = 473 nm) and the light output was measured by a power meter (Thorlabs) with a photodetector (Thorlabs) attached. Measurements were taken of fiber length of 1 - 10 cm, radii of curvature 1/8, 1/4, 3/8, 1/4, and 5/8 inch at a bending angle of 0° and 90°.

### Femtosecond laser micromachining

The fiber was immersed in dichloromethane for two minutes to remove the sacrificial outer PC layer before the micromachining process. The femtosecond(fs) laser used in the fabrication is a Ti: Sapphire NIR-fs pulsed laser (Coherent Libra series) with 800 nm emission wavelength, ~100 fs pulse width, and 3mm-beam-waist linear polarized Gaussian beam. A Nikon dry objective (20×, NA=0.4) was used to focus the laser beam on the fiber. The size of the focal area, *D*, is estimated at about 1.7 μm in diameter via *D*=2*λf*/π*w*_0_, where λ is the wavelength of the laser in the medium, f is the focal length of the lens, and w 0 is the input beam waist. The fiber was then mounted in an assembled motorized stage with a three-axis translation movement (ASI LX-4000 and Newport UTM50CC) and one-axis rotation around fiber axial direction (Thorlabs PRM1Z8). The fiber was fixed under tension to avoid wobbling during the fabrication. Since the laser beam only evaporates the material in the focal area, microchannel can be formed at any selected location by scanning the focused laser beam layer-by-layer from the surface. The average power used in the fabrication was 0.5 mW with the laser repetition frequency of 100 Hz. The scanning speed was 0.4 mm/s and the overlapping between each trace was around 1 μm. A reflected light microscopic system using the same objective was also implemented to monitor the fabrication in realtime. To characterize exposed microfluidic windows, 1 μl each of four different diluted food color dye (red, green, blue and copper, Wilton) was injected to separate microfluidic channels in the fiber through tubing connected to a standard precision injection apparatus (100 nl/s, NanoFil Syringe and UMP-3 Syringe pump, World Precision Instruments) while the fiber probe was embedded in 0.6% agarose gel (see details of tubing connection in Multifunctional Neuro Probes Assembly).

### Light scattering pattern measurement

The fiber was immersed in a drop of fluorescein solution (1% Uranine, Carolina Biological Supply Company) and excited by a 473 nm laser via butt coupling. The image was taken by an optical microscopy and the excitation light was filtered.

### Simulation of the exposed waveguide

The simulation was run by COMSOL Multiphysics© 5.5 equipped with frequency-domain electromagnetic wave solver on a workstation. Due to our limited computing power, two-dimensional simulation was preformed and it is enough to provide a qualitative evaluation. The dimension and geometry were determined by the microscopic image of the fabricated fiber. The waveguide has 11.5 μm thick PC core and 5 μm thick PMMA cladding. The exposed window is 20 μm x 20 μm wide and 7.5 μm depth with ~15° tapering. The side and bottom of the exposed window are roughened (R_a_=0.5 μm and 1μm for side and bottom respectively) to mimic the actual surface morphology of a femtosecond laser processed surface. The waveguide core is illuminated evenly with a 473 nm light and the corresponding refractive index of different layers are 1.6023 (PC),1.4976 (PMMA) and 1.3361 (water).

### Multifunctional Neuro Probes Assembly

For depth-dependent fiber probes used in *Thy1-ChR2-YFP* mice, the fiber probe was first put into the fiber optical ferrule and affixed by a retaining compound (LOCTITE) after micromachining process. Then the ferrule top part was polished by optical polishing papers from the roughness of 30um to 1um. Next, for electrical connection with the electrodes embedded in the fiber probe, the electrodes were exposed manually at different locations along the fiber length by a razor blade and silver paint (SPI Supplies) was applied to the manually exposed site individually. Then copper wires were wrapped around the fiber probe and additional silver paint was applied for a better connection. The wires were soldered to the pin connectors (Sullins Connector Solutions) while a stainless steel wire was soldered as a ground wire. Besides, 5-min epoxy (Devcon) was applied to the electrical and optical interface for affixation and electrical insulation. In the last step, we applied medical epoxy on the fiber probe tip area to prevent double exposure of the electrode during the electrophysiology recording process.

For spatially expandable fiber-based neural probes assembly, the sacrificial layer of the fiber probe was removed by dichloromethane immersion. To build microfluidic interfaces, the hollow channel of the fiber probe was exposed manually and then the fiber probe was inserted into the ethylene vinyl acetate tubing (0.5mm inner diameter) with the help of a needle syringe, resulting in the perpendicular positions of fiber probe and tubing with fiber probe’s microfluidic exposure site placed in the tubing center. Then 5-min epoxy was applied around the connection site to prevent leakage during microfluidic infusion to the fiber probe through the tubing. The electrical and optical interface connection was established via the methods described above. After the soldering process, fiber probes were inserted to the 5-channel scaffolding fiber manually and followed by the affixation process of the whole device with 5-min epoxy.

After the devices are fabricated, we will test their electrical, optical, and chemical functions before implant surgery to ensure all of the implanted devices are functional in the acute and long-term experiments.

### Surgical Procedure

All animal procedures were approved by Virginia Tech Institutional Animal Care and Use Committee and Institutional Biosafety Committee and were carried out in accordance with the National Institutes of Health Guide for the Care and Use of Laboratory Animals. Five to nine weeks old male wild type C57BL/6J mice (Jackson Laboratory) were received from the Jackson Laboratory (Bar Harbor, ME) and allowed to acclimate for at least 3 days before enrolling them in experiments. *Thy1-ChR2-YFP* mice were bred in our lab (Jackson Laboratory). Mice had access to food and water ad libitum and were kept in a facility maintained for 12 hours light/dark cycle starting at 7 AM. Male C57BL/6J mice were set up on a stereotaxic apparatus (David Kopf Instruments) and 1-3.5% isoflurane was induced to animals via nose cone during all procedures for anesthesia. To expose the scalp, a small incision was made on the skin along the midline then a small craniotomy was made with a dental drill. Then the assembled fiber probe was lowered using a micropositioner with respect to the Mouse Brain Atlas while the ground stainless steel wire was soldered to a miniaturized screw (J.I.Morris) on the skull. Finally, the whole exposed skull area was fully covered by a layer of Metabond (C&B METABOND; Parkell) and dental cement.

For surgeries with spatially expandable fiber probes used in spike and burst suppression recording, we first positioned the scaffolding fiber to the coordinates relative to bregma of −1.8 mm AP, 1.5 mm ML, −1 mm DV and affixed the scaffold using Metabond. Then the functional fiber probes were further inserted for 1 mm and the final coordinates of the five individual probes are (−1.8 mm AP, 1.3 mm AL, −2 mm DV), (−2 mm AP, 1.5 mm ML, −2 mm DV), (−1.8 mm AP, 1.5 mm ML, −2 mm DV), (−1.8 mm AP, 1.7 mm ML, −2 mm DV), and (−1.6 mm AP, 1.5 mm ML, −2 mm DV).

For surgeries with depth-dependent fiber probes in *Thy1-ChR2-YFP* mice, the coordinates relative to bregma used were −2 mm anteroposterior (AP); 1 mm mediolateral (ML); −3.3 mm dorsoventral (DV, four electrodes at −1 mm, −1.5 mm, −2.5 mm and −3.2 mm, respectively). For surgeries with drug intervention on *Thy1-ChR2-YFP* mice, the coordinates relative to bregma used were −2 mm AP, 1.5 mm ML, −3.2 mm DV.

For depth-dependent fiber probes used in seizure study with the straight implant, the coordinates relative to bregma used were −2 mm AP, 2 mm ML, −4.5mm DV with four electrodes at −1 mm, −1.5 mm, −2.5 mm and - 4.4 mm, respectively. For depth-dependent fiber probes used in seizure study at a 30° angled implant, the coordinates relative to bregma used were −2 mm AP, 1.5 mm ML (at the brain surface) and the distance of each exposed electrode from the brain surface was 0.5 mm (CTX), 1.3 mm (CA1), 2.2 mm (CA3) and 4.8 mm (AMG). The location of the electrode in the brain was verified in the coronal brain slices (50 μm thickness) after completion of the recording. The brain slices were stained for cell nuclei using DAPI to visualize clearly the damage caused by electrode implants (**Supplementary Figure 14**).

For surgeries with spatially expandable fiber probes used in *Thy1-ChR2-YFP* mice, we first lowered the scaffold to the coordinates of −1.5 mm AP, 0.5 mm ML and −2.5 mm DV relative to bregma. Then the scaffold was affixed to the scalp via Metabond while the fiber branches were further inserted into the brain for 1.5 mm after the Metabond was completely dry. Dental cement was used to affix the whole device above the skull. And the coordinates of the five electrodes are (−1.53 mm AP, 0.25 mm ML, −4 mm DV), (−1.75 mm AP, 0.53 mm ML, −4 mm DV), (−1.5 mm AP, 0.5 mm ML and −4 mm DV), (−1.47 mm AP, 0.75 mm ML, −4 mm DV), and (−1.25 mm AP, 0.47 mm ML, −4 mm DV).

For surgeries with spatially expandable fiber probes used in *Thy1-ChR2-YFP* mice to induce the seizure-like aftercharges, the coordinates of the scaffolding fiber is −2 mm AP, 1.5 mm ML, −0.6 mm DV and the functional fiber probes were further inserted for 0.5 mm (−2 mm AP, 1.4 mm ML, −1.1 DV) to target cortex region, 1 mm (−2.15 mm AP, 1.5 mm ML, −1.6 DV) to target CA1 region rostrally, 1 mm (−1.85 mm AP, 1.5 mm ML, −1.6 mm DV) to target CA1 region caudally, and 1.5 mm (−2 mm AP, 1.75 mm ML, −2.1 mm DV) to target CA3 region.

For surgeries with spatially expandable fiber probes used in seizure study, the coordinates for the scaffold relative to bregma used here are −2 mm AP, 1.5 mm ML, −1 mm DV. The lengths of inserted functional fibers beyond the bottom of the scaffolding fiber were 0.5 mm (CA1), 1 mm (CA3), 2 mm (thalamus) and 2 mm (thalamus). And the coordinates of the four electrodes are (−2.1 mm AP, 1.5 mm ML, −1.5 mm DV), (−2 mm AP, 1.7 mm ML, −2 mm DV), (−1.6 AP, 1.5 mm ML, −3 mm DV), and (−2 mm AP, 1.1 mm ML, −3 mm DV).

### Viral Infection

Theiler’s murine encephalomyelitis virus (TMEV) was generously provided by Drs. Karen S. Wilcox and Robert S. Fujinami, University of Utah. Mice were infected with TMEV as previously reported^61^. Mice were anesthetized using 3% isoflurane and kept under anesthesia using 2-3% isoflurane during the entire infection procedure. The injection area in the right hemisphere was disinfected with 70% ethanol and the injection site is pinpointed slightly medial to the equidistant point on the imaginary line connecting the eye and the ear. A William’s collar syringe which contains a plastic jacket on the needle to expose only 2 mm of the needle was used for injection to restrict the injection within the somatosensory cortex. About 20-25 μl of Daniels strain of TMEV solution containing 300,000-375,000 plaque forming units (PFU) of the virus was injected intracortically by inserting the needle at a 90° angle to the skull. The needle was kept in place for at least 1 minute before retracting slowly to prevent leakage.

### In Vivo Electrophysiology

For studies involving single-unit LFP investigation in wild type mice and for the measurement of optically-induced electrical activities in *Thy1-ChR2-YFP* mice, a 32 Channel Neurophysiology System (Tucker-Davis Technologies, TDT) was connected to the headpins of the implanted device after animals recovered from surgeries. For optical stimulation, the fiber probe was connected to the DSPP laser as mentioned above. During optical stimulation, a laser pulse with 5 ms pulse was used and the frequencies used were 10, 20, and 100 Hz. Stimulation was delivered in 0.6 s stimulation epochs every 5 seconds. For the drug intervention experiments, the concentration of 0.1 mM CNQX (Tocris) solutions in PBS was prepared and 2.5 μl of which was injected into the brain through fiber probe’s hollow channel using NanoFil Syringe and UMP-3 Syringe pump system at the speed of 80 nl/s.

### Video-electroencephalography (vEEG)

After 4-7 days of recovery from the surgical procedure, mice were infected with TMEV as described above and enrolled for continuous vEEG recordings between 2 and 8 days post-TMEV infection. The MP160 data acquisition system and AcqKnowledge 5.0 software from BIOPAC Systems, Inc. were used to record electroencephalograms. The corresponding behavior of each mouse was recorded using Media Recorder 4.0 software (Noldus Information Technology) and M1065-L network camera (Axis communications). Mice were connected to EEG100C differential amplifiers (BIOPAC) using custom-adapted six-channel cable connectors (363-000 and 363-441/6, Plastics1) and six-channel rotating commutators (SL6C/SB, Plastics1). All the cables and electrical components were sufficiently shielded to minimize electrical noise. Food and water were freely accessible to mice during the entire vEEG recording. Video and EEG recordings were automatically synchronized using the Observer XT 14.1 software from Noldus Information Technology. EEG signals were bandpass-filtered (high pass filter: 0.5 Hz, low pass filter: 100 Hz), amplified, and digitized at a sampling frequency of 500 Hz.

### Data Analysis

Data analysis was performed with Matlab (The Mathworks) and custom scripts were used to sort neural spikes and analyze the local field potentials (LFPs). The activities were digitally filtered from 0.5-300 Hz and 300-5000 Hz to present single spike resolutions and LFP resolution. The spike sorting algorithm was implemented by filtering out individual spikes using standard deviation dependent threshold, reducing the dimensionality of the data via principal component analysis (PCA) and using unsupervised learning algorithms to separate out the clusters, such as K-means clustering. A custom Matlab script was also written to create spectrograms to visually support the analysis of the LFPs in both the time domain and the frequency domain. The EEG and video recordings were reviewed manually for the detection of seizures. Electrographic seizures were defined as fast rhythmic spikes or sharp-wave discharges with amplitudes at least two times higher than baseline and lasting for at least 5 s. EEG signals during the ictal period typically start with low amplitude spikes that gradually increase in amplitude during seizure progression and then gradually decrease in amplitude and frequency at the end of the seizure. Suppression of EEG baseline also commonly occurs following convulsive seizures corresponding to behavioral arrest in mice. By verifying these gradual rhythmic changes in EEG signals and the suppression of basal EEG activity, seizures were identified and artifacts were excluded from the analysis.

### Immunohistochemistry

Animals were implanted with a stainless-steel probe or a Fiber F1/F2 for 4 weeks and then anesthetized with a ketamine (20 mg/mL)/xylazine (2 mg/mL) solution and transcardially perfused with phosphate-buffered saline (PBS) (Fisher BP661-10) followed by 4% paraformaldehyde (PFA) (Electron Microscopy Sciences Cat. #15714-S) in PBS. Upon extraction, the brain was kept in 4% PFA overnight at 4□°C and then placed into PBS containing 0.02% sodium azide (Sigma S8032). The brain was serially sectioned into 50 μm transverse slices on a Campden Instruments 5100mz vibratome. All slices were blocked for 1hr at room temperature in blocking solution that contained 50% goat serum (Millipore S26-100ML) and 0.01% Triton X-100 (Sigma T9284) in PBS containing 0.02% sodium azide. Slices were then subsequently blocked in 8.3% Affinipure Fab Fragment goat anti-mouse IgG solution in 1X PBS. After blocking, slices were incubated with primary antibodies diluted in blocking solution without Triton X-100 overnight at room temperature. Primary antibodies used included chicken anti-GFAP (abcam Cat. #ab4674, 1:1000), mouse anti-NeuN (Millipore Cat. #MAB377, 1:100), and rabbit anti-Iba1 (Wako Cat. #019-19741, 1:500). Following primary incubation, slices were washed six times with PBS for 10 minutes at room temperature with agitation. Secondary antibodies diluted in blocking solution without Triton X-100 were added at room temperature for 1 hr. Secondary antibodies used included goat anti-chicken Alexa Fluor 488 (Jackson ImmunoResearch Cat. #103-545-155, 1:500) goat anti-mouse Alexa Fluor 647 (Jackson ImmunoResearch Cat. #115-605-062, 1:500) and goat anti-rabbit Cy3 (Jackson ImmunoResearch Cat. #111-165-144, 1:500). Slices were then washed six times with PBS for 10 min at room temperature with agitation. Slices were mounted on glass slides with VECTASHIELD Antifade Mounting Medium with DAPI (Vector Laboratories Cat. #H-1200). Optical sections were acquired using a A1R Nikon laser scanning microscope using a Plan Apo 20X/N.A.0.75 air objective. Quantification of the data was performed using NIS-Elements (Nikon Instruments Inc.). To compensate for variance in lesion size, the puncture wound area was measured, and circular inner and outer regions of interest (ROI) were generated to form an annulus with an area normalized to all slices (A = π(R^2^ - r^2^)). Neuron density was then calculated within the normalized area by counting NeuN labeled cell bodies using spot detection (Nikon Elements). Area analysis of IBA1 and GFAP labeled cells was performed by creating binary layers of the optical sections using the threshold tool and quantified using the measurement tool (Nikon Elements). Projection images were created using ImageJ (NIH) software.

### Statistics

Parametric statistical tests were used if the data were sufficiently normal distributed and variance within groups was sufficiently similar. Experimental designs with two comparison groups were analyzed by a two-tailed unpaired t-test and designs with more than two comparison groups were analyzed using analysis of variance (ANOVA). The effects of CNQX on electrophysiological recordings were analyzed by repeated measure one-way ANOVA and Tukey’s multiple comparisons test. The power of electrographic recording obtained from different brain regions was compared by two-way ANOVA and Tukey’s multiple comparisons test. The difference between the two groups was considered statistically significant with a p-value of less than 0.05. Significance of tissue reaction to chronically implanted multimodal Fiber F2 and conventional stainless steel probes after four-week implantation was determined by the student’s t-test. All tests were performed using GraphPad Prism 8.0.

### Data availability

The data that support the findings of this study are available from the corresponding author upon reasonable request. The source data underlying Figs. 5e, i, 6e, f, and g and Supplementary Figs. 12d, 13e, f, and g are provided as a Source Data file.

### Code availability

The MATLAB scripts for analysis are available from the corresponding author upon reasonable request.

