## Supplementary information for "Spatially expandable fiber-based probes as a multifunctional deep brain interface"

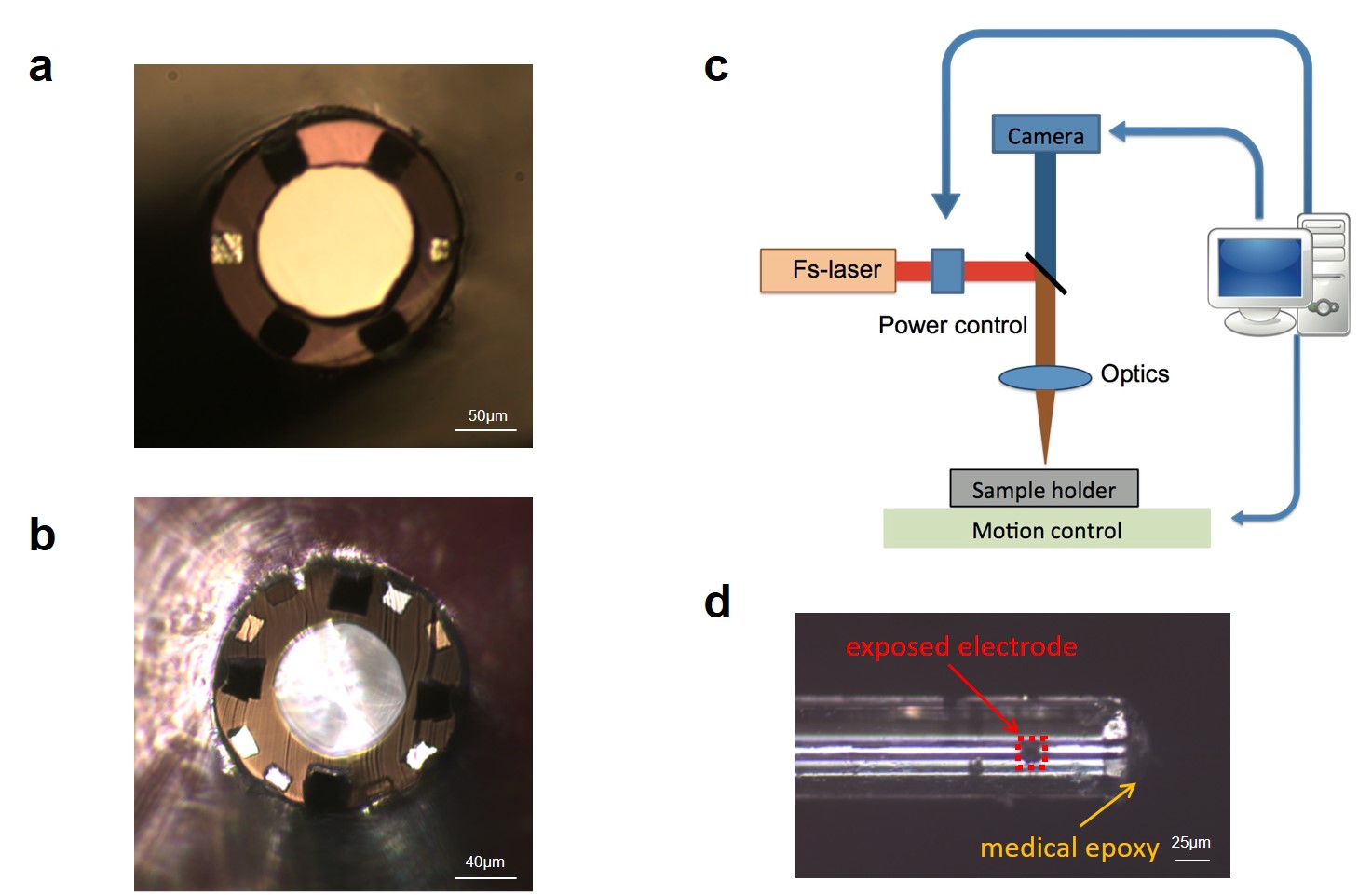

**Supplementary Figure 1: Depth-dependent fiber probes (Fiber S1-2)** (**a**) A cross-sectional image of the Fiber S1. (**b**) A cross-sectional image of the Fiber S2. (**c**) A detailed illustration of the femtosecond laser micromachining platform. (**d**) A microscope image of the exposed electrodes.

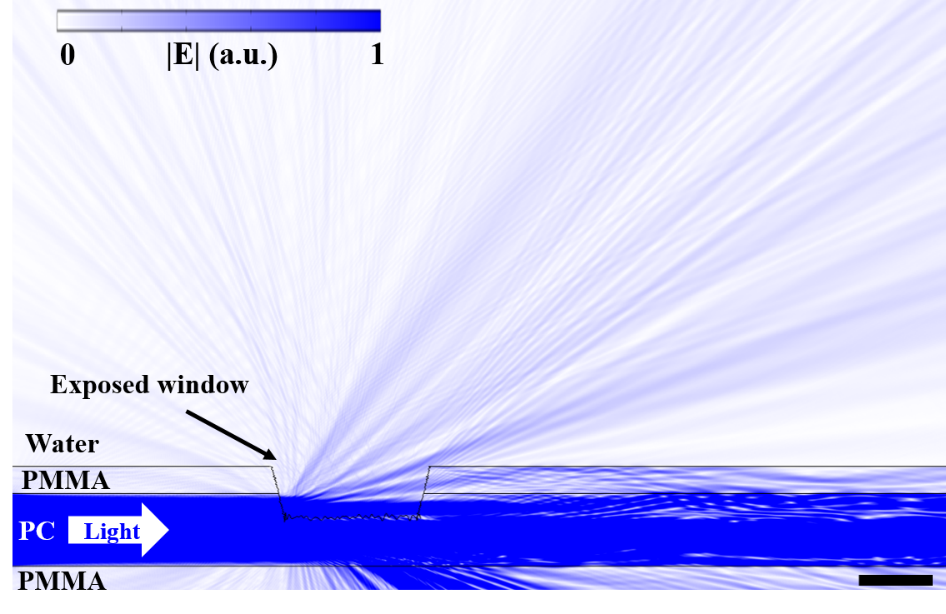

**Supplementary Figure 2: 2D simulation of the scattered light intensity from a femtosecond-laser exposed waveguide.**

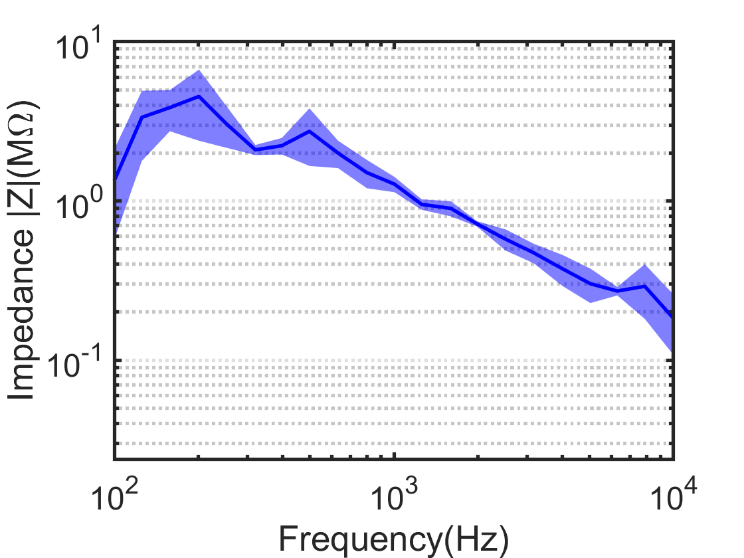

**Supplementary Figure 3: EIS measurement of the exposed electrode of Fiber F3.** The impedance spectrum of this exposed electrode of Fiber F3 confirmed the successful exposure of the electrode after femtosecond laser micromachining.

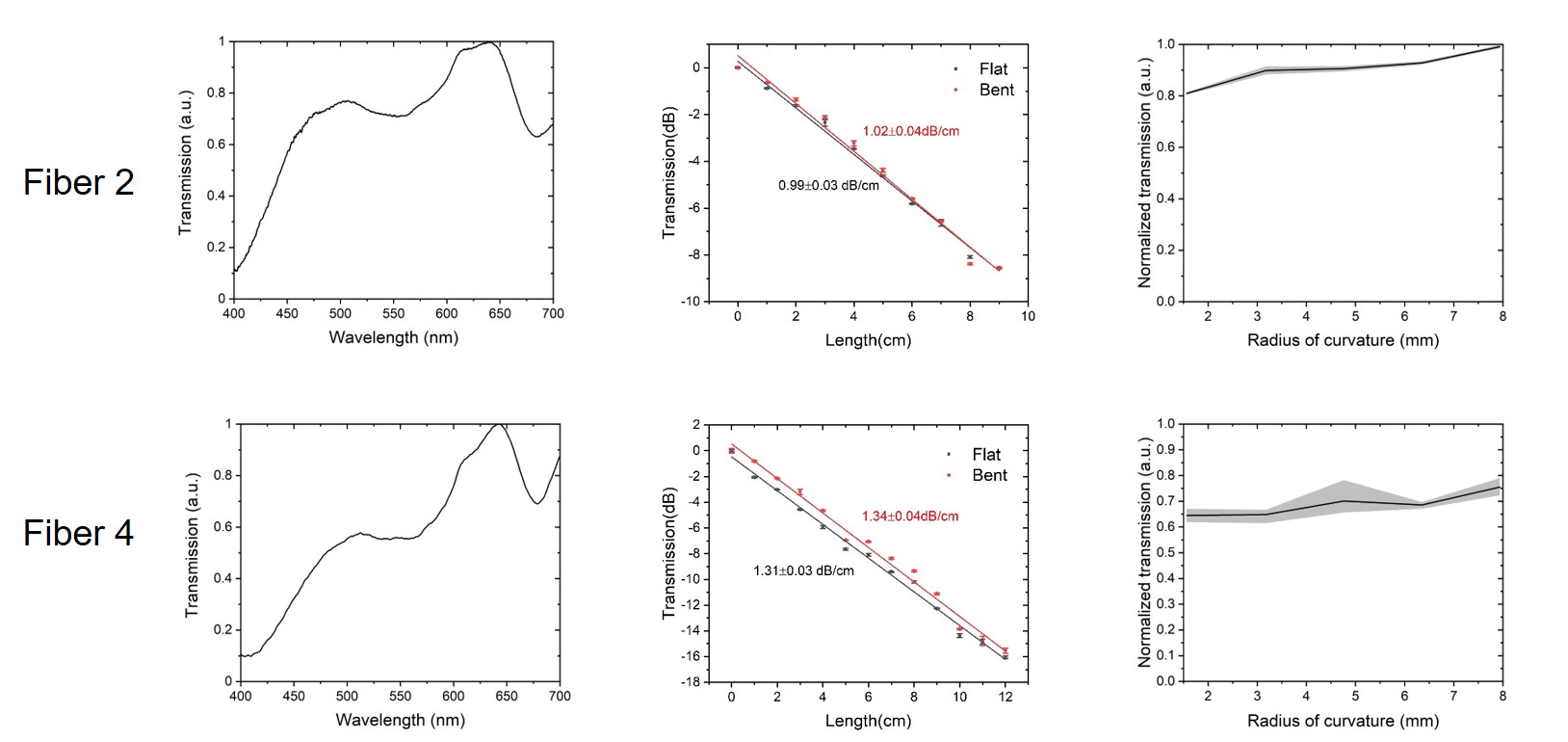

**Supplementary Figure 4: Characterization of optical properties of Fiber F2 and Fiber F4.** The optical transmission spectrum was obtained at a wavelength range of 400 – 700 nm, showing broad transmission across the visible range (normalized to the maximum value). The transmission loss measured using cut back method is 0.99±0.03 dB/cm (F2, R^2^=0.99308, n=3) and 1.31±0.03 dB/cm (F4, R^2^=0.99274, n=3) at a wavelength of 473 nm. And the bending test at 90º angle and radii of curvature of 1.6-7.9 mm shows no significant bending loss under these deformation conditions.

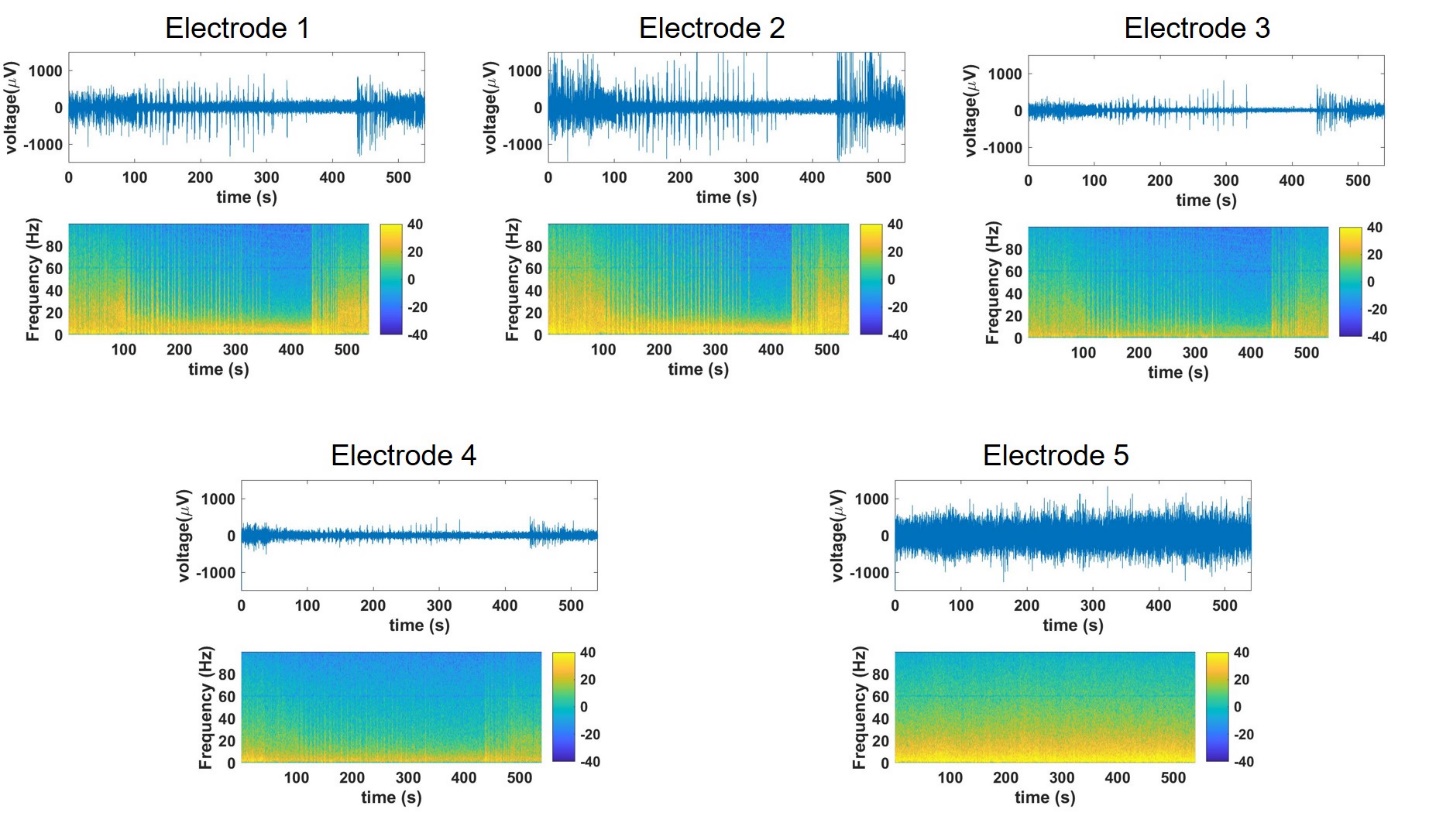

**Supplementary Figure 5: LFP recordings from spatially expandable fiber probes implanted in the hippocampus region of the wild type mouse brain with altered isoflurane levels during recording (0.5-2% v/v).** Electrodes 1-4 showed similar results.

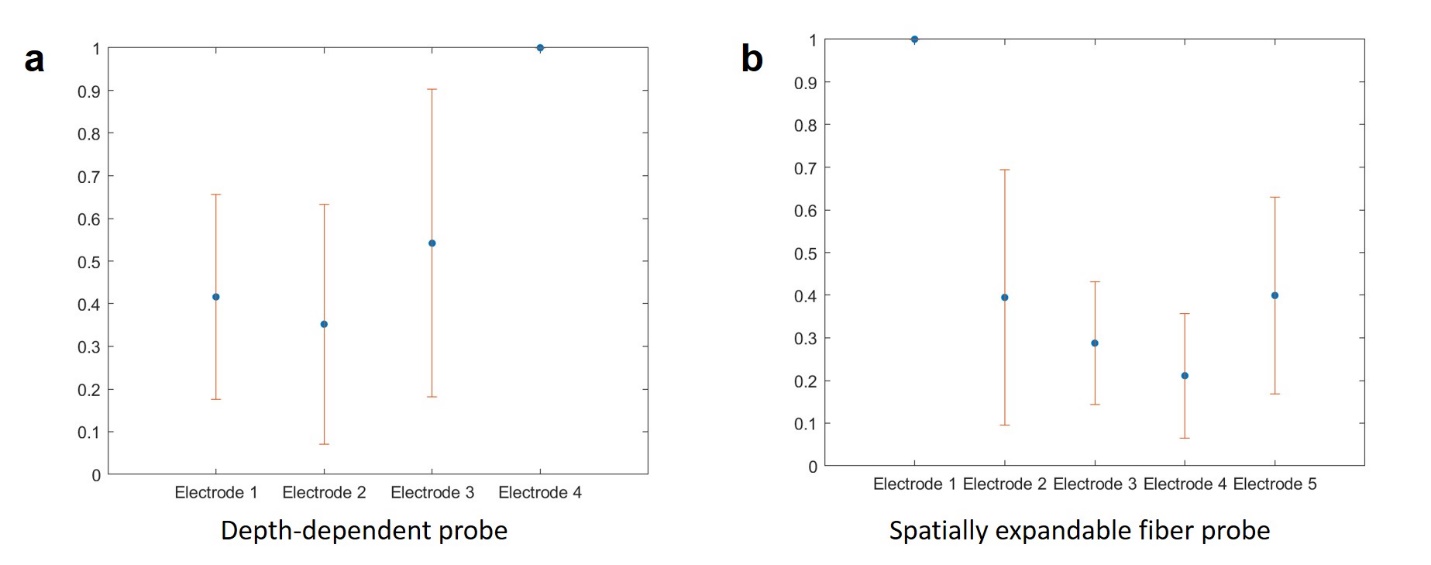

**Supplementary Figure 6: Normalization of the recording results from depth-dependent probes and spatially expandable fiber probes.** (**a**) Normalized optically stimulated results from depth-dependent probes indicate that the electrode four which was the closet to the optical stimulation site detected the highest signal in average. (**b**) Normalized optically stimulated results from spatially expandable fiber probes indicate that the electrode one which was the closet to the optical stimulation site detected the highest signal in average. All error bars in the figure represent the standard deviation.

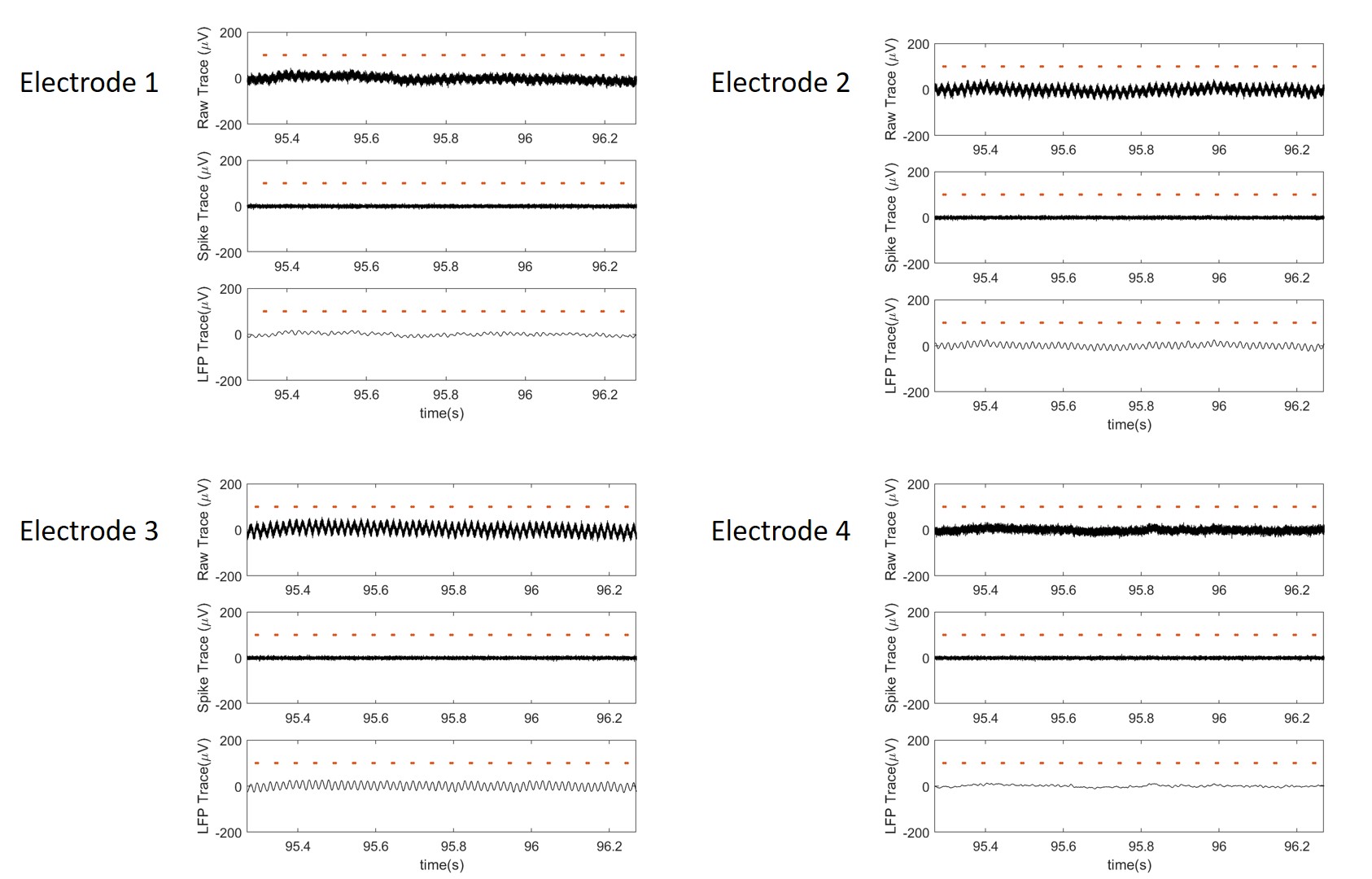

**Supplementary Figure 7: Recordings from a perfused brain with optical stimulation of 10 Hz by the depth-dependent fiber probes.** No optically evoked signals recorded from these four electrodes confirmed the fact that the recording results we collected in live animals were not artifacts.

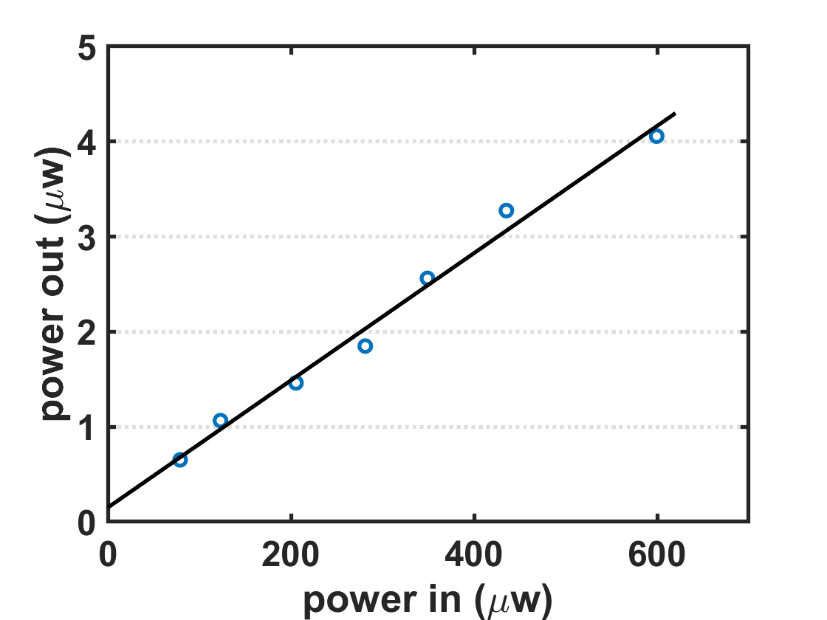

**Supplementary Figure 8: Light efficiency measurement of a single exposed optical window.**

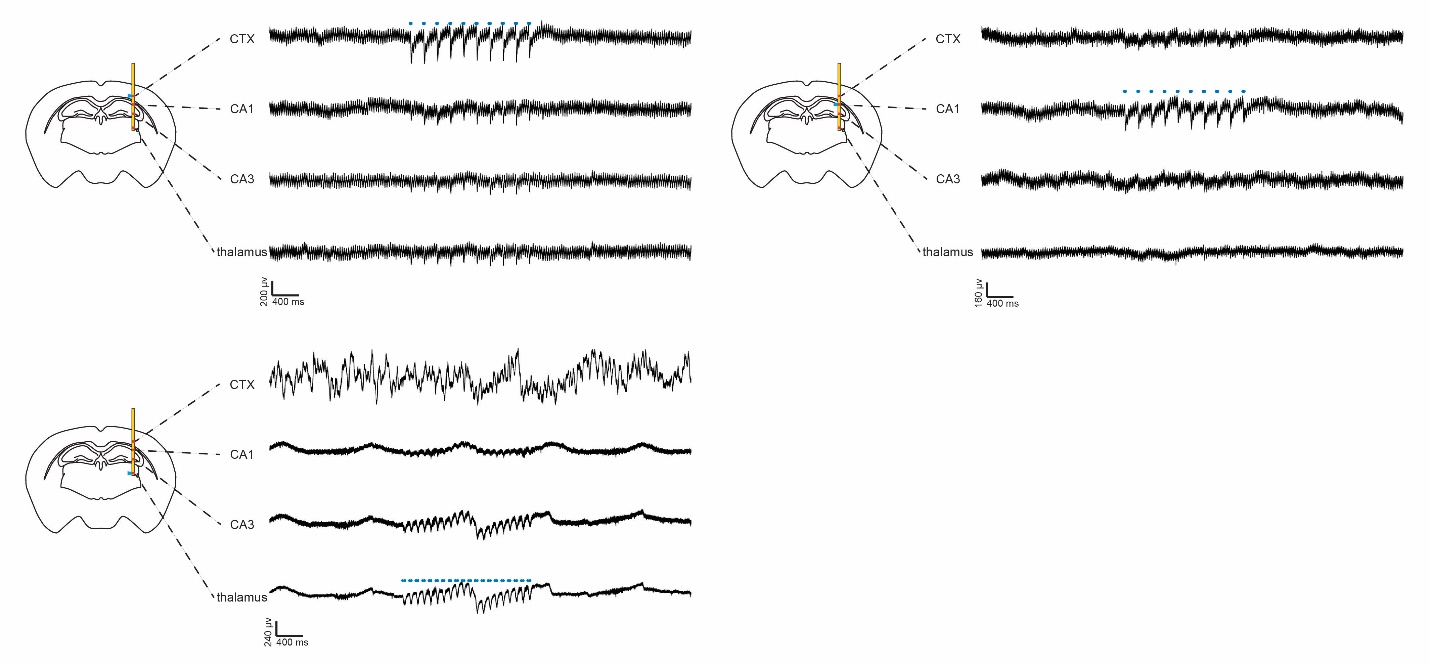

**Supplementary Figure 9: Simultaneous optical stimulation and electrical recording in Thy1 mouse using the depth-dependent fiber probes**. The four electrodes are located at the cortex, CA1, CA3, and thalamus regions, and light is delivered at the side of the fiber adjacent to the electrodes targeting cortex, CA1, and thalamus regions, respectively. The blue dashed lines indicate the laser pulses.

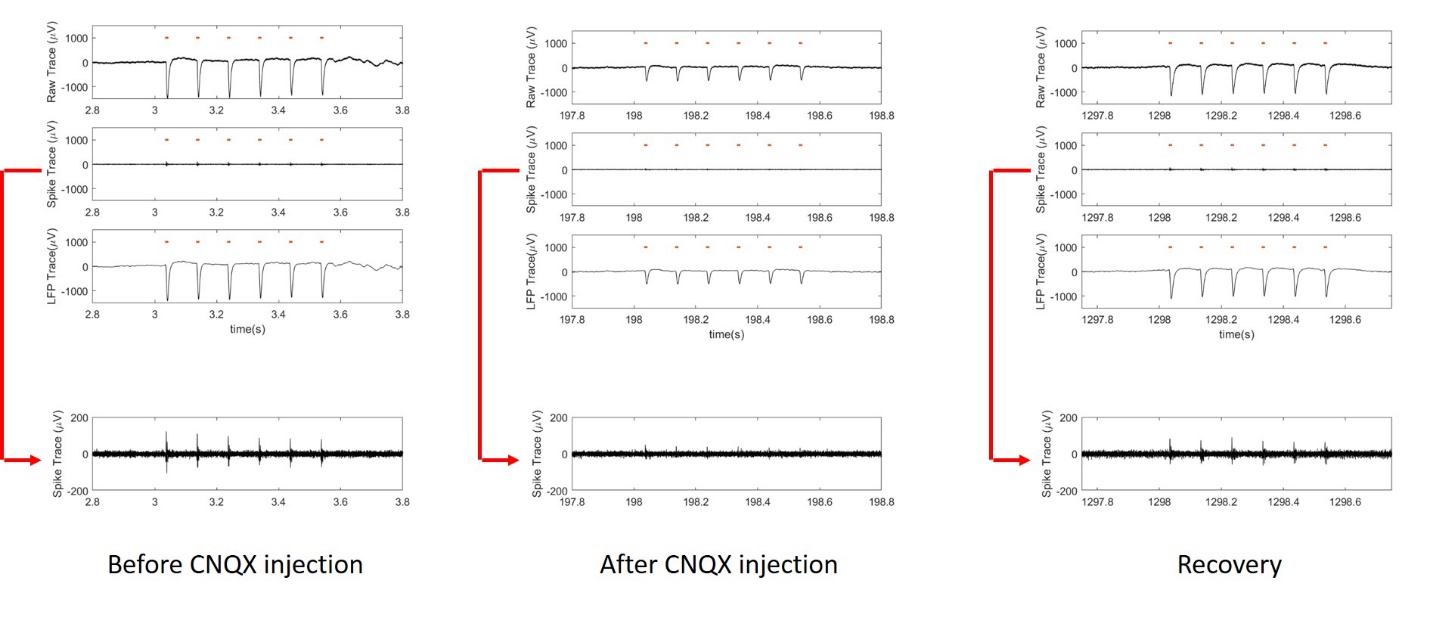

**Supplementary Figure 10: Unfiltered, Spike-filtered, and LFP filtered recording results before, during and after CNQX administration in a transgenic *Thy1-ChR2-YFP* mouse brain.** We can observe that the spike-filtered recording had no optically evoked signal immediately after CNQX injection while the raw data still presented the optically evoked signal in the whole time, which indicates that CNQX only affected neurons close to the microfluidic channel while the neighboring neurons that were not in contact with CNQX still responded to laser pulses.

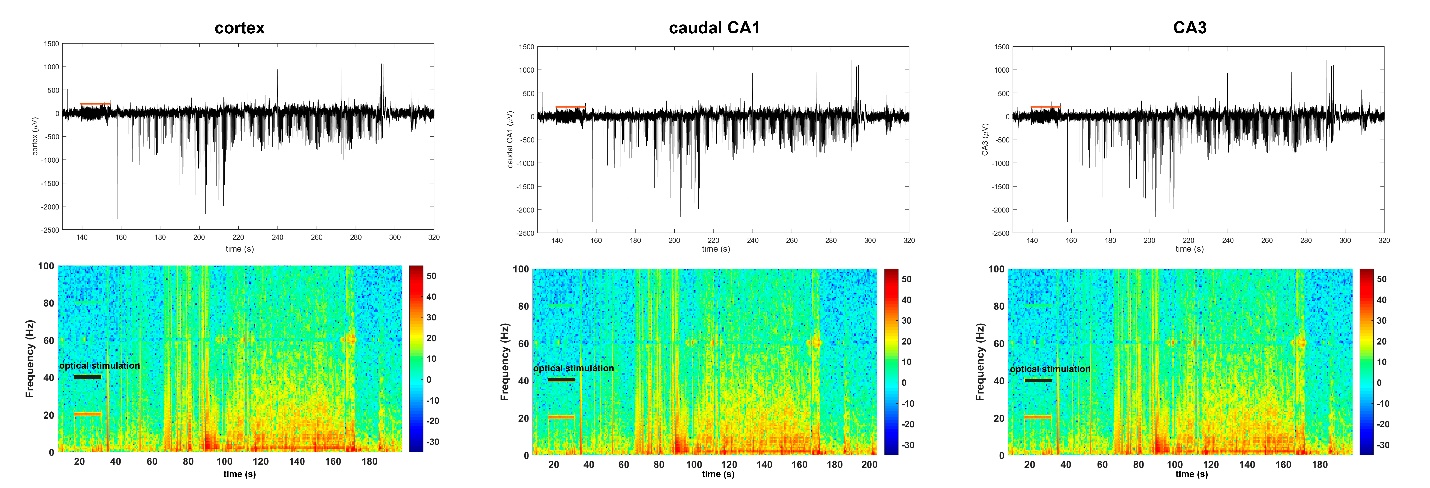

**Supplementary Figure 11: Similar brain activity of the seizure-like afterdischarges detected by other three brain regions.**

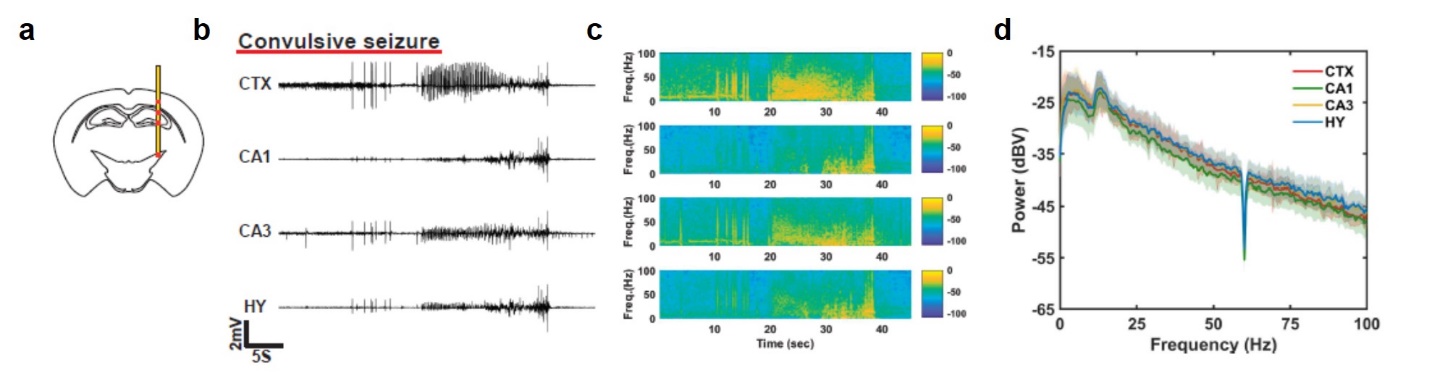

**Supplementary Figure 12: Convulsive seizure detected by depth-dependent fiber probe (straight implant).** (**a**). Diagram of coronal section of a mouse brain shows the location of depth-dependent fiber probes in cortex (CTX), CA1 and CA3 regions of hippocampus, and hypothalamus (HY) as indicated by red marks along with the probe assembly (straight implant). (**b-c**). Representative traces of electrographic recordings from CTX, CA1, CA3, and HY during a non-convulsive seizure and their corresponding frequency distribution. (**d**). Power spectrum density plot for convulsive seizures shows no difference in mean power of signal recorded from different regions.

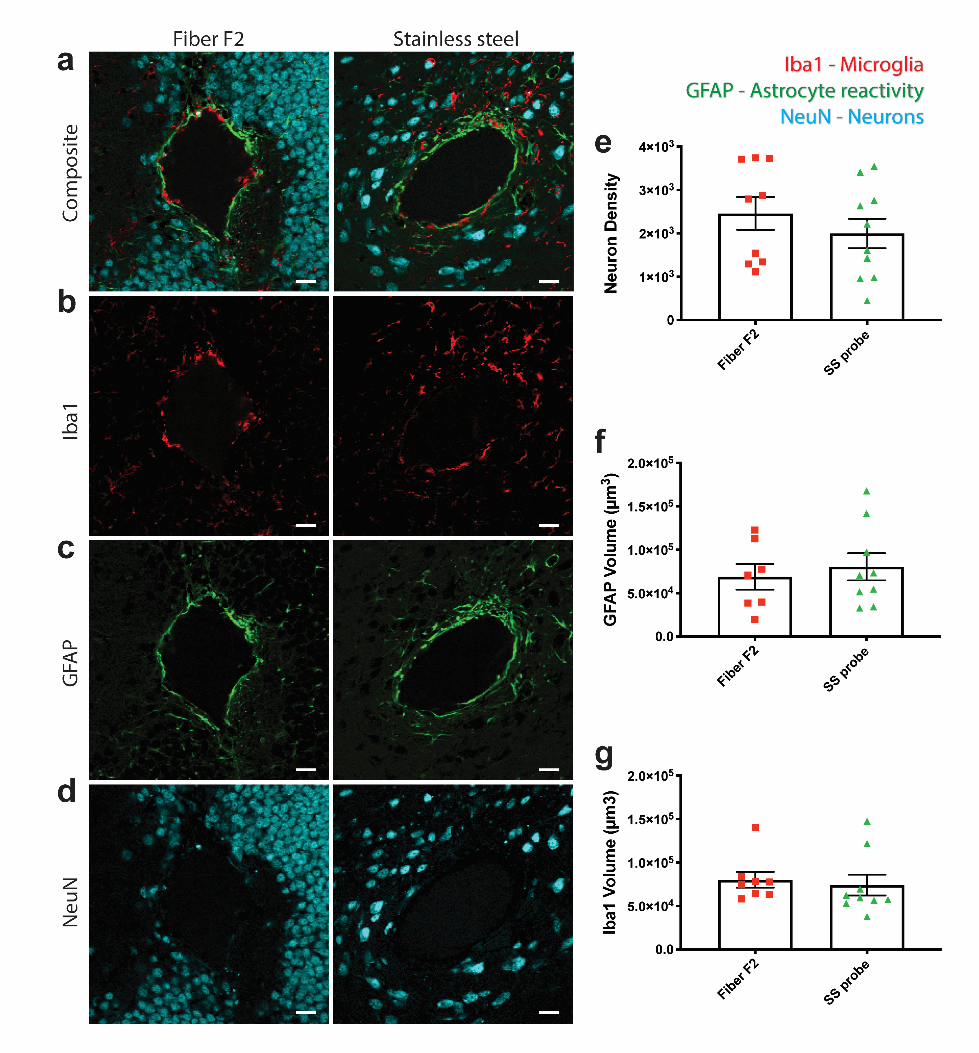

**Supplementary Figure 13: Immunohistochemical comparison of tissue reaction to chronically implanted multimodal Fiber F2 and conventional stainless steel probes after four-week implantation.** (**a−d**) Single confocal optical sections were taken at electrode implantation site. Neurons were labeled with NeuN (cyan), astrocytes with GFAP (green), and microglia with Iba1 (Red). (**d,e**) Neuron density, calculated by counting NeuN labeled neurons, was not significantly different between the groups. (**c,f**) Astrocyte and microglia (**b,g**) reactivity, measured as the volume of GFAP- or Iba1-positive cells respectively, was not significantly different between groups. Significance was determined by student’s t-test. Error bars on bar graphs reflect the standard deviation. (Scale bar: 40 μm)

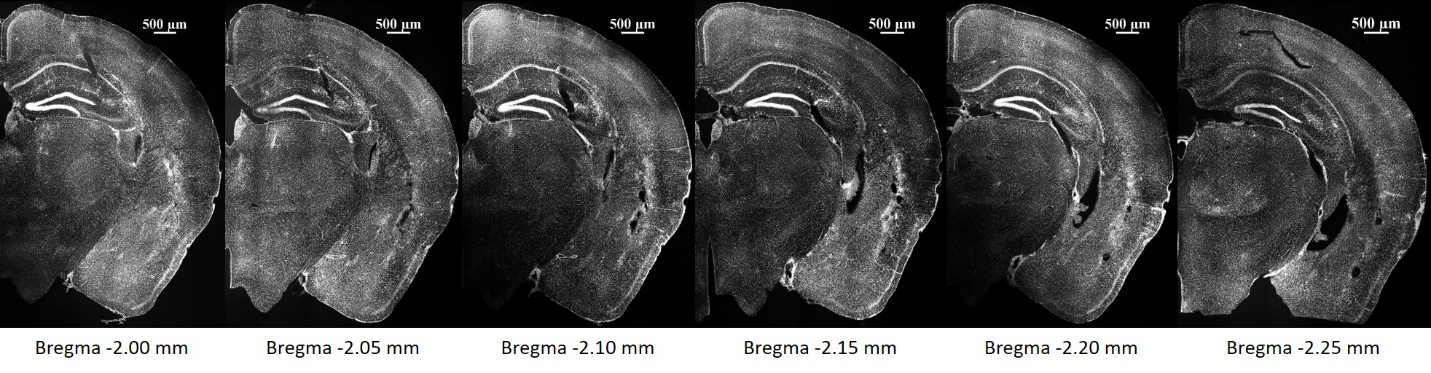

**Supplementary Figure 14: Location of depth-dependent fiber probe implanted angularly to the brain surface for electrophysiological recording.** Mice brains collected at the end of electrographic recordings were sliced (50 µm thickness) and stained with DAPI to visualize cell nuclei. Serial coronal sections from 2.00 to 2.25 mm posterior to bregma are shown from left to right. Diagonal cut mark seen in each slice indicates location of the electrode implanted to target cortex, CA1 and CA3 regions of hippocampus, and amygdala.
